# Defining T cell receptor repertoires using nanovial-based affinity and functional screening

**DOI:** 10.1101/2023.01.17.524440

**Authors:** Doyeon Koo, Zhiyuan Mao, Robert Dimatteo, Natalie Tsubamoto, Miyako Noguchi, Jami McLaughlin, Wendy Tran, Sohyung Lee, Donghui Cheng, Joseph de Rutte, Giselle Burton Sojo, Owen N. Witte, Dino Di Carlo

## Abstract

The ability to selectively bind to antigenic peptides and secrete cytokines can define populations of cells with therapeutic potential in emerging T cell receptor (TCR) immunotherapies. We leverage cavity-containing hydrogel microparticles, called nanovials, each coated with millions of peptide-major histocompatibility complex (pMHC) monomers to isolate antigen-reactive T cells. T cells are captured and activated by pMHCs and secrete cytokines on nanovials, allowing sorting based on both affinity and function. The TCRs of sorted cells on nanovials are sequenced, recovering paired αβ-chains using microfluidic emulsion-based single-cell sequencing. By labeling nanovials having different pMHCs with unique oligonucleotide-barcodes we could link TCR sequence to targets with 100% accuracy. We identified with high specificity an expanded repertoire of functional TCRs targeting viral antigens compared to standard techniques.

**One-sentence Summary:** Affinity and secretion-based screening of antigen-specific T cells using nanovials defines a functional TCR repertoire

## Main Text

In the future, engineered cell therapies will be a pillar of medicine along with molecular and genetic interventions. There have been encouraging successes in the use of engineered T cell-based therapies, including T cell receptor (TCR) immunotherapy in treating cancer. These approaches use endogenous signaling activity in T cells and rely on the recognition of cancer-associated antigens that are presented as peptides associated with major histocompatibility complex (MHC) on the surface of tumor cells (*1*). Engineered TCRs have demonstrated efficacy in treating multiple types of tumors including melanoma, sarcoma, and leukemia (*2, 3*).

One technical hurdle for developing effective TCR immunotherapy is to identify reactive TCRs that can recognize targets with sufficient affinity and potency to induce apoptosis or activate other immune cells by secreting communication factors, such as cytokines. T cells have one of the most diverse sequence repertoires (10^8^-10^20^) to respond to a wide variety of pathogens (*4, 5*). Recent single-cell screens have also highlighted an astonishingly high level of functional heterogeneity from T cells isolated from the same patient and bearing the same panel of surface markers, with only a small highly active subset of cells driving responses to immunological challenge (*6*–*8*). Current tools for enriching and screening cognate T cell populations rely mostly on TCR affinity or function, as defined by surface markers of cell activation and intracellular cytokine staining in fixed/dead cells. Peptide-MHC (pMHC) multimer (e.g. tetramer) staining is the conventional method to specifically label T cells with cognate TCRs benefiting from the avidity effect when four pMHC monomers are linked through a tetrameric streptavidin backbone (*9*). However, pMHC multimer staining does not take into account the functional stages of the T cells, and TCR affinity is not always correlated with activation or cytotoxicity (*9*). An alternative way to isolate reactive T cells is through surface activation markers (e.g. CD25, CD69, CD137). The selection of T cells based on activation biomarkers can be achieved without the knowledge of specific epitopes and the readouts of these markers are based largely on functional activation of the T cell rather than TCR affinity (*10, 11*). Despite improvement in these activation-based selection techniques and better choices of markers, some of the isolated T cells have been reported to be “bystander T cells’’, meaning they were not able to respond to antigens in a reconstructed experiment (*11*). As the size of the pool of antigens increases, the deconvolution of TCR-restricted targets can also become laborious (*10*–*12*). Our previous efforts led to the development of the “CLint-seq” technique, in which a reducible crosslinker was used to fix T cells that were stained intracellularly for cytokines as activation markers (*13*). The processed cells by CLint-seq can still be sequenced at the single-cell level and show highly specific recovery of functional TCRs. Since intracellular cytokine staining (ICS)-based techniques require fixation and permeabilization, it leads to less RNA recovery at lower quality and still requires deconvolution of target epitopes. An ideal technology would combine antigen-specific enrichment and activation-based screening to achieve both highly specific identification of functional TCR sequences along with the knowledge of their cognate target epitopes.

We recently reported an approach to confine cells in small nanoliter-volume cavities within hydrogel microparticles, which we call ‘nanovials’, and capture secreted molecules on the nanovial surfaces (*14, 15*). Here, we adapted the nanovial technology to achieve combined antigen-specific capture and functional activation-based high-throughput analysis and sorting of T cells based on secreted cytokines (Fig. 1). Each nanovial acts as both an artificial antigen-presenting cell that presents millions of pMHC molecules within the cavity to capture with high avidity and activate cells with cognate TCRs, and as a capture site for secreted cytokines. To recover TCR sequences in an epitope-specific manner, live cells on nanovials are sorted based on CD3 and CD8 expression and interferon-γ (IFN-γ) secretion, followed by single-cell sequencing to construct a single-cell TCR library with matching αβ-chains. Corresponding antigen-specific information is linked to each TCR sequence using an oligonucleotide feature barcode encoding the specific pMHC molecules on the nanovial. We demonstrate the ability to isolate thousands of rare antigen-reactive T cells and construct a TCR library with significantly expanded breadth and epitope-specific annotation compared to a library constructed from cells enriched using standard approaches that were tested in parallel.

**Figure 1.**
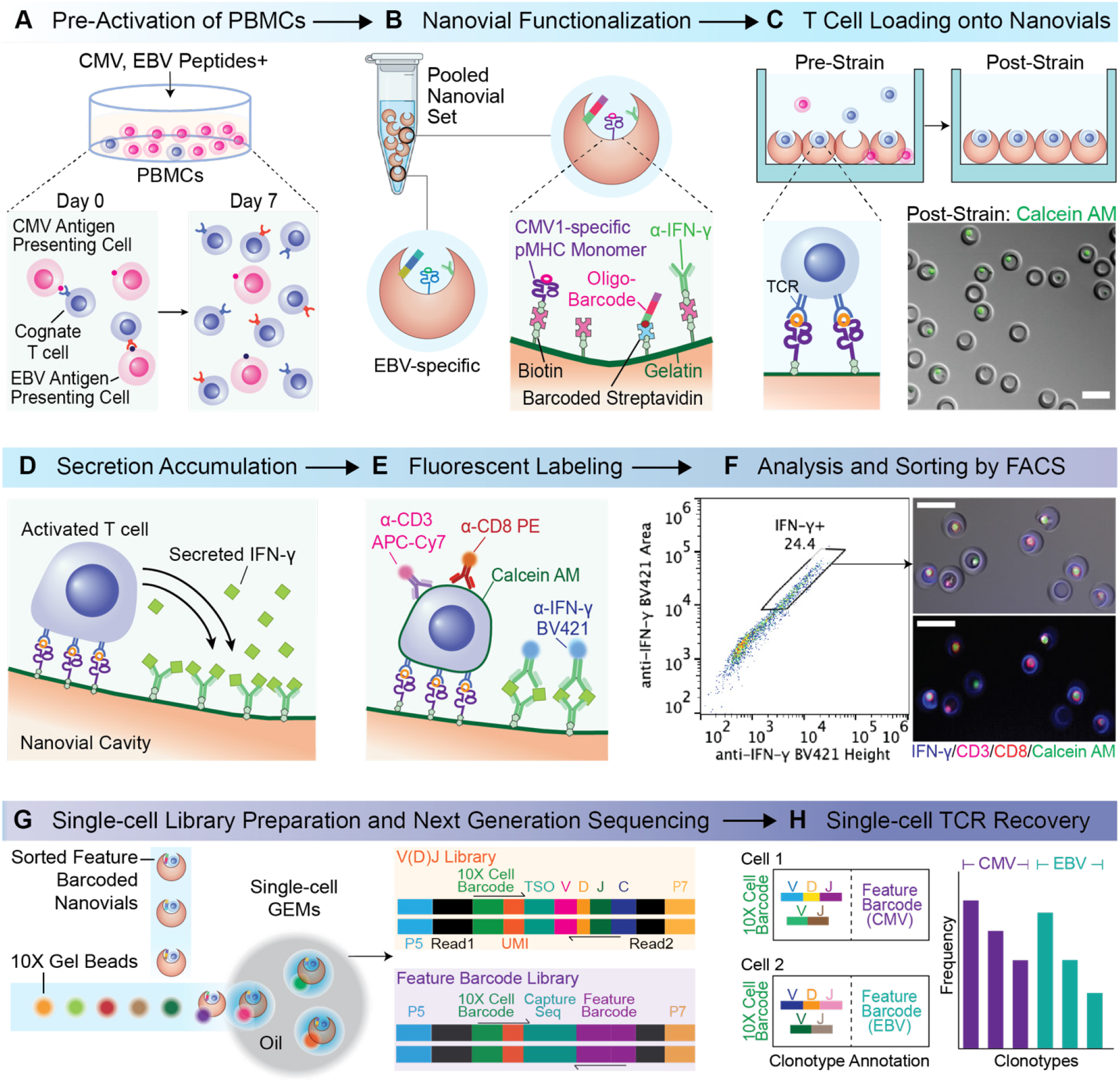
Overview of high-throughput analysis and isolation of antigen-specific T cells followed by recovery of a single-cell TCR library. (A) Pre-activation of PBMCs with cytomegalovirus (CMV) and Epstein-Barr virus (EBV) specific peptides for 7 days. (B) Functionalization of nanovials with cytokine capture antibodies, pMHC monomers and oligonucleotide barcodes via streptavidin-biotin chemistry. (C) Loading of cognate T cells into the cavities of nanovials in a well plate and removal of unbound cells using a cell strainer. (D) Activation of T cells for 3 hours and secreted cytokine capture in the cavity of nanovials. (E) Labeling of captured cytokines and cell surface markers with fluorescent detection antibodies. (F) Sorting of cells on nanovials based on viability, CD3/CD8 expression and secretion signal. (G) Compartmentalization of sorted population into droplets with a cell barcode bead in the 10X Chromium system for the construction of matched V(D)J and feature barcode (nanovial barcode) libraries. (H) Annotation of TCR clonotypes with corresponding epitopes by matching feature barcode. Scale bars represent 50 μm.

## Results

### Fabrication and functionalization of nanovials for T cells

Our first aim was to functionalize nanovials to capture T cells and three important inflammatory cytokines. We used a microfluidic device that generates uniform water-in-oil emulsions to create millions of monodisperse polyethylene glycol (PEG)-based nanovials with an inner cavity selectively coated with biotinylated gelatin (fig. S1A) (*14, 15*). To accommodate human T cells with diameters of ∼10 μm we fabricated uniform nanovials with an average outer diameter of 35 μm (CV = 5.1%) and an average cavity diameter of 21.2 μm (CV = 7.2%). By functionalizing the inner cavity with biotin during fabrication we could flexibly link multiple biotinylated antibodies or peptide-MHC (pMHC) monomers with epitopes of interest through streptavidin-biotin noncovalent interactions. For capture and secretion analysis of primary human T cells, irrespective of antigen targeting, we decorated nanovials with biotinylated anti-CD45 and cytokine capture antibodies against interferon-γ, tumor necrosis factor-α, and interleukin-2 (anti-IFN-γ, anti-TNF-α, anti-IL 2). At least a two order of magnitude dynamic range in detection of recombinant cytokines was observed when anti-CD45 capture antibodies and one 1:1 (140 nM each) or two cytokine secretion capture antibodies 1:1:1 (140 nM each) were used during functionalization (fig. S1B). All of these conditions allowed for T cell loading (fig. S2) as well as signal capture from recombinant cytokines down to 10 ng/mL (fig. S1B). Cell loading followed Poisson loading statistics with an optimum observed when 1.6 cells per nanovial were seeded (fig. S2A) with anti-CD45 used to capture cells (fig. S2B), and was largely independent of the presence of additional cytokine capture antibodies (fig. S2C). To capture antigen-specific T cells, nanovials were decorated with pMHC monomer along with cytokine capture antibodies. We saw a linear increase in loaded pMHC monomer signal up to concentrations of 80 µg/mL. Assuming all biotinylated monomers bound to nanovials we estimated there to be a maximum of ∼160 million pMHC monomers per nanovial (fig. S1C). In order to maintain sites for capture antibodies targeting secreted cytokines we limited the pMHC concentration to 20 µg/mL (∼30,000 pMHCs/μm^2^) for future experiments unless otherwise stated. For comparison, ∼200,000 pMHC molecules (∼640 pMHCs/μm^2^) were reported to be on the surface of T cells (*16*). The ability to load high surface densities of pMHC monomer in the nanovials is expected to result in higher avidity binding to TCRs than alternative techniques.

Having validated recombinant cytokine assays and T cell loading on nanovials we wanted to test the processes to accumulate and detect secretions from single T cells bound to nanovials using flow cytometric analysis. We developed assays for three secreted cytokines (ΙFN-γ, ΤΝF-α, IL-2) using nanovials coated with the respective individual cytokine capture antibody and anti-CD45. After loading primary human T cells onto nanovials, cells were activated non-specifically with phorbol 12-myristate 13-acetate (PMA) and ionomycin for 3 hours. Following fluorescent staining of captured cytokines, we used a standard cell sorter, the SONY SH800S, to sort nanovials with each cytokine secretion signal based on fluorescence peak area and height values (Supplementary Note1). By also gating on a cell viability dye, such as calcein AM, we were able to simultaneously measure secretions and viability of individual cells on nanovials, improving the selective sorting of functional cells (fig. S3A). We gated and sorted populations into low, medium, or high secretors based on the area vs. height plot, recovering viable cells with different levels of TNF-α and IFN-γ secretion as reflected in fluorescence microscopy images (fig. S3B). We also evaluated crosstalk between nanovials by co-culturing cell-loaded nanovials and fluorescently labeled (AlexaFluor 488) nanovials without cells and found that less than 0.04% of test nanovials without cells appeared in the positive secretion gate during the 3 hour activation period (fig. S3C). Presumably, secreted cytokines accumulate at higher concentrations locally but when cytokines diffuse or are advected to neighboring cavities, the concentration is diluted substantially leading to a reduced signal.

### Capture of antigen-reactive T cells with pMHC-labeled nanovials

We hypothesized that nanovials coated with pMHC and cytokine capture antibodies could be used for antigen-specific capture, TCR-specific activation, and detection of secreting cytokines. Transitioning from using anti-CD45, pMHC-functionalized nanovials were applied for the selection of antigen-reactive T cells. We first analyzed the specificity of nanovials in selectively binding antigen-specific T cells using human peripheral blood mononuclear cells (PBMCs) transduced with 1G4 TCR targeting NY-ESO-1, a clinically studied cancer-specific antigen (*3, 17*). Truncated nerve growth factor receptor (NGFR) was used as the co-transduction marker for the presence of 1G4 TCR. PBMCs transduced with and expressing 1G4 bound specifically to nanovials labeled with pMHC monomer containing HLA-A*02:01 restricted NY-ESO-1 C9V peptide (SLLMWITQV) (20 µg/ml) (Fig. 2A). Live cells occupied ∼17% of nanovials, and 93.9% had NGFR expression. To further clarify if the interaction was specific, we increased the NY-ESO-1 pMHC concentration used to functionalize nanovials. A corresponding increase in the amount of antigen-specific T cells loaded was observed in a dose-dependent manner, while the binding of untransduced PBMCs was low and independent of pMHC concentration (Fig. 2B, fig. S4A).

**Figure 2.**
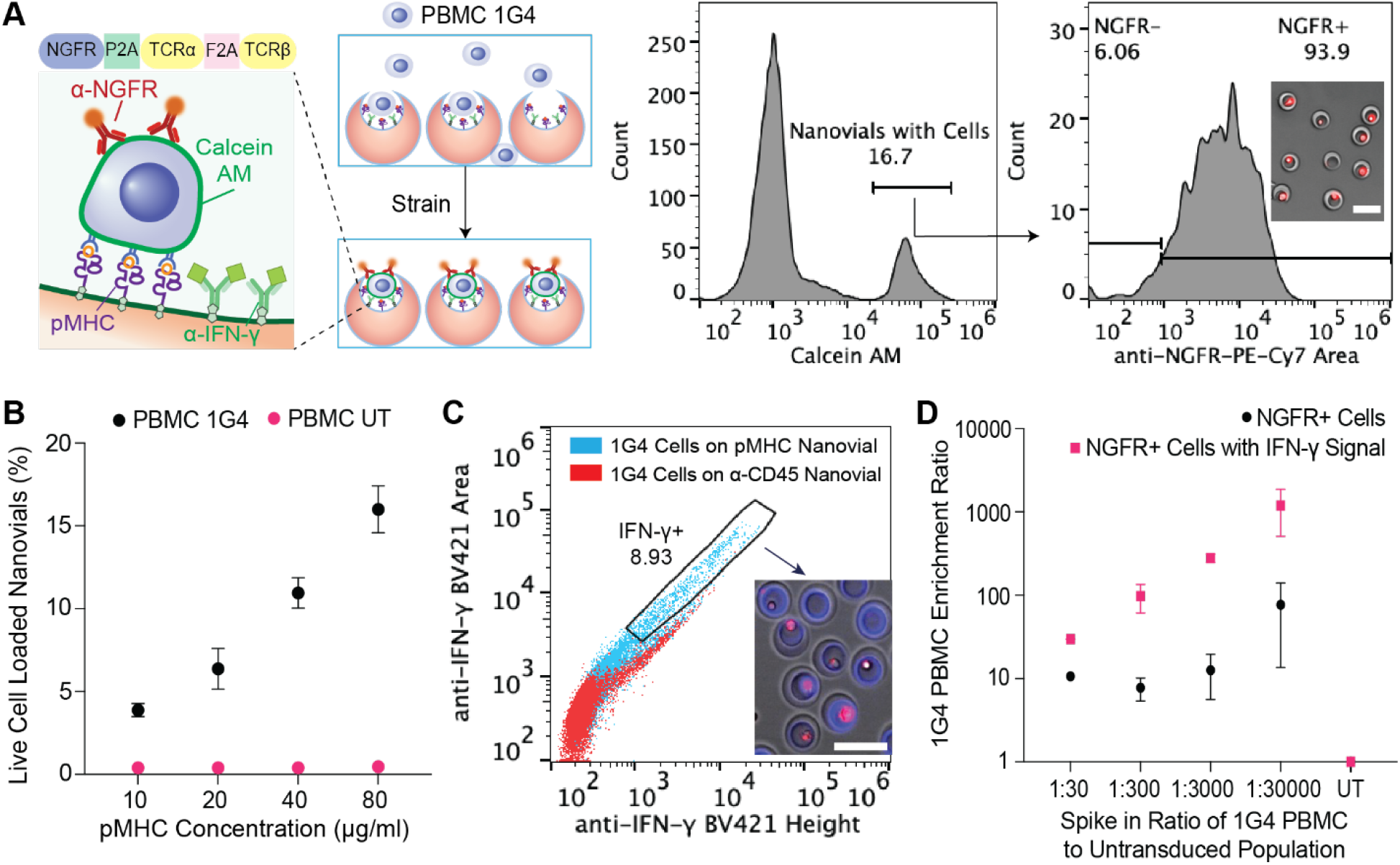
Detection of antigen-specific T cells on HLA-A*02:01 restricted NY-ESO-1 pMHC labeled nanovials. (A) PBMCs transduced with 1G4 TCRs are captured onto NY-ESO-1 pMHC labeled nanovials with ∼94% of bound cells staining positive for anti-NGFR. (B) The fraction of nanovials containing live cells plotted as a function of pMHC concentration for 1G4-transduced (black dots) or untransduced PBMCs (magenta dots). (C) Flow cytometry plots of IFN-γ secretion induced by nanovials at 3 hours for 1G4-transduced PBMCs loaded onto anti-CD45 labeled nanovials (red dots) or NY-ESO-1 pMHC labeled nanovials (cyan dots). Secreting cells on pMHC-labeled nanovials sorted from the gated area are shown. (D) Enrichment ratios for 1G4-transduced PBMCs diluted into untransduced PBMCs at different ratios (1:30, 1:300, 1:3000, 1:30000), when considering all bound cells (black dots) or bound cells with IFN-γ signal (magenta dots). Scale bars represent 50 μm.

To investigate whether pMHCs on nanovials can specifically trigger activation and secretion from engaged antigen-specific T cells, 1G4-transduced cells were each loaded onto nanovials labeled with pMHC monomers or anti-CD45 antibodies. Cells loaded on control anti-CD45 labeled nanovials (red dots) had low levels of IFN-γ signal which was mostly associated with non-specific staining of cells, while cells on NY-ESO-1 pMHC-labeled nanovials (cyan dots) clearly secreted IFN-γ as early as 3 hours after loading (fig. S4B), yielding 1-2 orders of magnitude higher fluorescence intensity observed at larger area:height ratios (Fig. 2C).

Because of the expected rarity of antigen-reactive T cells, we then investigated the sensitivity of pMHC-coated nanovials to specifically recover cognate T cells from 1G4-transduced PBMCs diluted into untransduced PBMCs at different ratios and loaded onto NY-ESO-1 pMHC labeled nanovials. We assessed enrichment with both binding alone (CD3^+^/CD8^+^/NGFR^+^) and binding combined with positive secretion signal (CD3^+^/CD8^+^/NGFR^+^/IFN-γ^High^) (fig. S5). An enrichment ratio at each spiking ratio was calculated by dividing the final fraction of NGFR positive population when considering all bound cells or bound cells with IFN-γ signal by the initial fraction of NGFR positive population at each dilution. We were able to detect 1G4 transduced cells up to 1:30000 dilution (enrichment ratio of 1000), where combining binding with cytokine secretion signal increases the specificity of detection by 15-fold (Fig. 2D).

For some therapeutic workflows enrichment and regrowth of rare antigen-reactive populations are required. To assess proliferation of antigen-specific T cells after isolation, 1G4-transduced PBMCs enriched on NY-ESO-1 pMHC-coated nanovials were first sorted and detached using collagenase D. Cells expanded in culture over 5 days, with 90% of the expanded population expressing the 1G4 TCR, showing enrichment and continued growth of the antigen-specific T cell population (fig. S4C).

### T cells with low affinity TCRs are isolated effectively by pMHC-coated nanovials

Since the 1G4 TCR has high affinity to NY-ESO-1 pMHC, we questioned whether increased avidity of pMHCs coating the nanovial cavity would prove advantageous in recovering TCRs with various affinities. Human PBMCs were transduced with five previously identified TCRs (3A1, 1G4, 4A2, 5G6, 9D2) targeting the same HLA-A*02:01 restricted NY-ESO-1 C9V peptide. The relative affinities of these TCRs were assessed using fluorescent pMHC dextramer binding (*17*). We compared the recovery efficiency and purity of antigen-specific cells by nanovial capture, nanovial capture gated on IFN-γ secretion, and dual-color tetramer staining. pMHC-labeled nanovials recovered more antigen-specific T cells than dual-color tetramer, especially when cells possessed low-affinity TCRs (4A2, 5G6, 9D2), suggesting that the high avidity of the nanovial surface is sufficient to allow binding of T cells independent of their affinities (Fig. 3A). When gating on the presence of IFN-γ secretion, recovery efficiency was reduced, since not all cells secrete, but recovery purity of the isolated cells approached 100% (Fig. 3A). The purity of detected cells was defined as the fraction of NGFR^+^ cells from CD3^+^/CD8^+^ cells on nanovials with or without IFN-γ secretion signal, or fraction of NGFR^+^ cells from dual-tetramer^+^ cells. When PBMCs transduced with each TCR were spiked into untransduced PBMCs at different ratios, antigen-reactive T cells were enriched across all five TCRs to a similar level up to 1:100 dilution (Fig. 3B).

**Figure 3.**
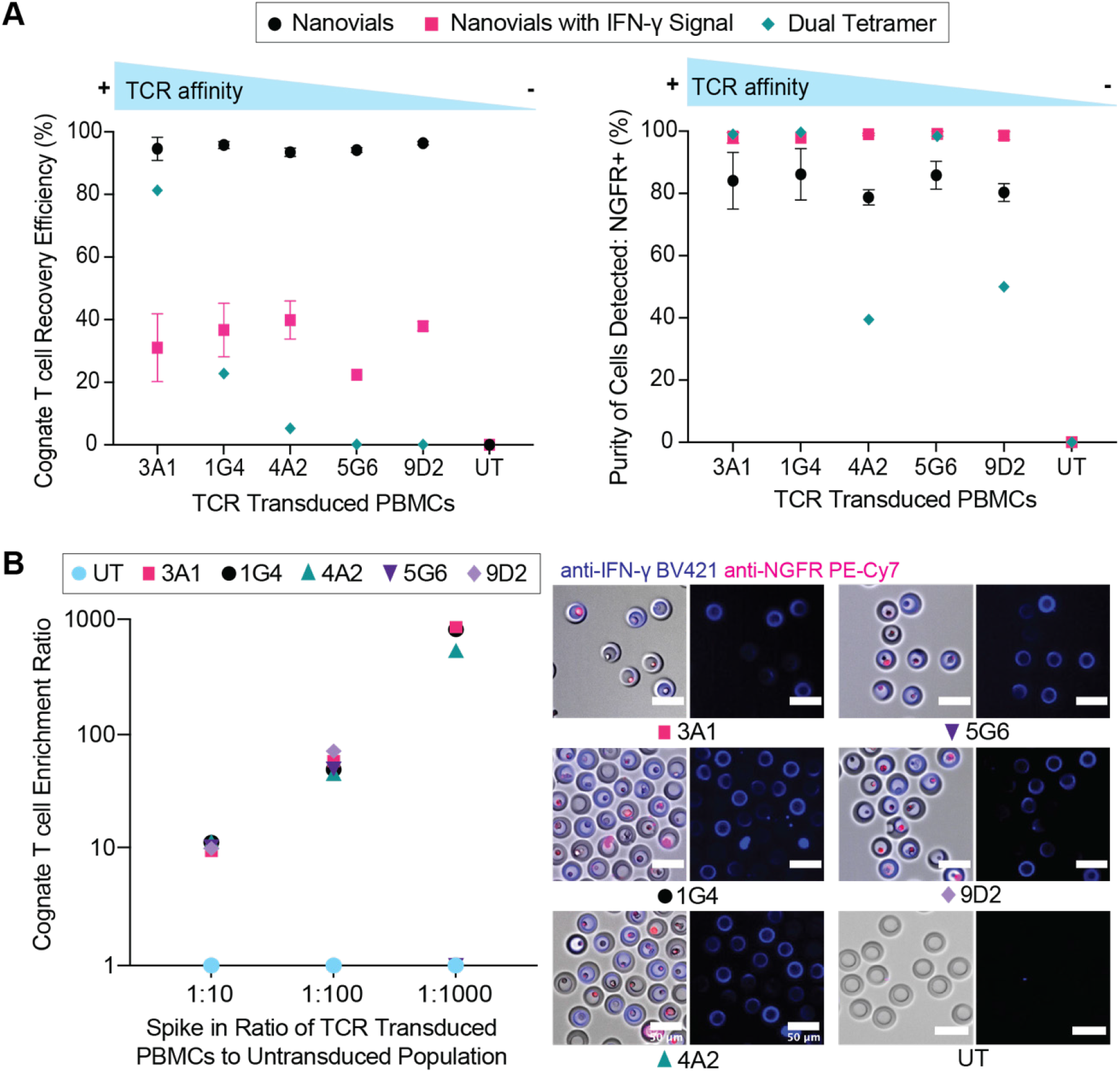
Recovery of cognate T cells with various affinities to HLA-A*02:01 restricted NY-ESO-1 pMHC. (A) The detection efficiency and purity of sorted populations of cells based on binding to nanovials (circles), nanovials with IFN-γ secretion (squares) and labeling with dual-color tetramers (Dual Tetramer, diamonds) as a function of TCR. (B) The enrichment ratio of cells transduced with TCRs possessing different affinity as a function of spiked fraction (1:10, 1:100, 1:1000). Microscopy images of sorted cells with IFN-γ secretion signal for each transduced TCR. Scale bars represent 50 μm.

### pMHC-nanovials yield a broader repertoire of TCRs compared to tetramers

Following successful isolation of rare antigen-specific T cells with low-affinity TCRs in model systems we hypothesized that sorting based on a combination of affinity and cytokine secretion using nanovials would recover a more diverse repertoire of highly functional TCRs compared to existing techniques. Healthy donor PBMCs pre-activated with a pool of previously reported HLA-A*02:01 restricted peptides from cytomegalovirus (CMV) and Epstein Barr virus (EBV) targeting CMV pp65 (CMV1, NLVPMVATV), CMV IE-1 (CMV2, VLEETSVML), and EBV BMLF1 (EBV, GLCTLVAML) were isolated using three different methods: affinity and secretion based sorting using nanovials, affinity-based sorting using a CMV pp65-specific tetramer, or activation-based sorting using CD137 as the surface marker (Fig. 4A). We multiplexed the detection of antigen-specific T cells by loading cells onto a pool of barcoded nanovials labeled with three HLA-A*02:01 restricted pMHCs targeting each antigen (CMV1, CMV2, EBV) and a corresponding oligonucleotide barcode. Approximately 6000 CD3^+^CD8^+^ cells on nanovials associated with IFN-γ secretion signal and 800 CMV1-specific pMHC tetramer+ cells were identified and sorted from the entire sample (fig. S6, Fig. 4B). To have an equivalent starting cell number for single-cell sequencing, 6000 cells were sorted based on gating for above background levels of the CD137 activation marker (fig. S6).

**Figure 4.**
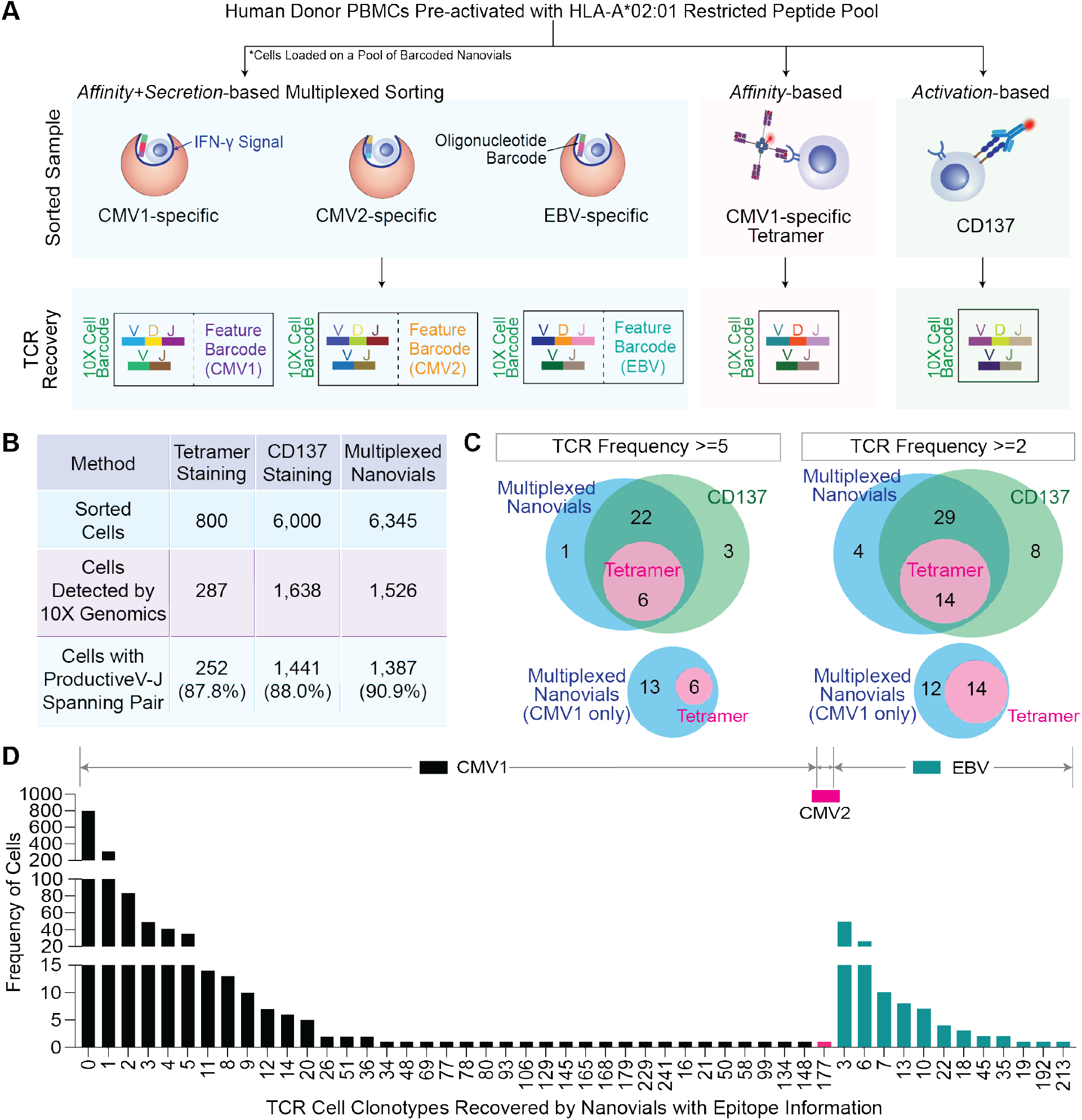
Sorting of antigen-specific cells and recovery of single-cell TCR clonotypes using nanovial, tetramer, or CD137 approaches. (A) TCR discovery workflow and matching epitope deconvolution using nanovials along with comparison techniques. For nanovials, each TCR was matched with a corresponding oligonucleotide barcode sequence reflecting pMHC information. (B) Summary of single-cell TCR αβ sequencing results. Sorted cells refers to the number of cells gated and sorted as positive (see fig. S6 for detailed gates). (C) A representative Venn diagram of recovered TCR clonotypes from three approaches with a frequency≥5 (left) or ≥2 (right). (D) Distribution of cell clonotypes recovered by nanovials color coded with matching target pMHC information: 32 CMV1 (black), 1 CMV2 (magenta), 12 EBV (green) specific clonotypes.

TCRs were recovered from sorted cells using the 10X Genomics Chromium platform (Fig. 4B). Notably, the cells on nanovials were introduced directly into the system to maintain the connection between a nanovial with a feature barcode oligonucleotide tag and the attached T cell. Nanovials did not interfere with the gene sequence recovery resulting in the highest fraction of cells with a productive V-J spanning pair (90.9%) compared to tetramer (87.8%) and CD137 samples (88%). We compiled a list of high-frequency TCR clonotypes (frequency≥5) detected by the three methods. Since a few clonotypes contained multiple alpha or beta chains, we recombined them into separate TCR sequences with each permutation of alpha and beta chains. In total, we retrieved 32 unique TCR pairs with frequency≥5: 6 TCRs overlapped among the three methods, 28 overlapped between the nanovial and CD137 approaches, and one unique TCR was detected with nanovials (Fig. 4C). A larger number of unique TCR sequences were detected with a less stringent cutoff of frequency≥2, where the overlapping number of TCRs between nanovial and CD137 techniques increased from 28 to 43, suggesting additional rarer TCRs were also discovered.

### Barcoded nanovials reveal epitope information during T cell isolation

Unlike workflows using CD137, which require laborious deconvolution to uncover the target epitopes from a peptide pool that match specific TCR sequences, pMHC-barcoded and multiplexed nanovials reveal epitope information during cognate T cell isolation. Using the 10X Chromium system, TCR sequence information of each cell was linked to the nanovial pMHC feature barcode, resulting in the recovery of each TCR with matching target epitope information. A >90% frequency of the pMHC barcode identified the dominant epitope for each TCR (fig. S7). The distribution of clonotype frequency with the corresponding epitope is represented in Fig. 4D for the nanovial workflow. We detected 32 CMV1(CMV-pp65), 1 CMV2(CMV-IE1) and 12 EBV-reactive clonotypes and 96.4% of sequenced cells with productive V(D)J spanning pairs had matching epitope information.

To address the antigen-specific reactivity of 32 TCR sequences retrieved by the three methods (nanovial, tetramer, CD137) with a frequency≥5, candidates were re-expressed via electroporation into Jurkat-NFAT-GFP cells, in which GFP expression can be induced upon TCR recognition. Murine constant regions were used for both TCR alpha and beta chains to prevent mispairing with endogenous TCRs. Engineered Jurkat cells were then co-cultured with K562 cells expressing HLA-A*02:01 (K562-A2) as antigen-presenting cells along with exogenously added peptides. Activation of the Jurkat cells was determined by flow cytometry, gating on % of the CD8^+^/murineTCRβ^+^ population with GFP signal above background. From the smaller pool of CMV1-specific TCRs, the nanovial workflow yielded 6 more reactive TCRs compared to CMV1 pMHC tetramer labeling (Fig. 5A). Out of the 29 possible TCR combinations identified with nanovials, 17 were found to be reactive upon re-expression (11 for CMV1 and 6 for EBV) (Fig. 5A). Notably, some of these TCRs were from cells in which sequencing yielded multiple alpha and/or beta chains (denoted with connecting line and an asterisk, Fig 5A). When collapsing these related TCR clonotypes to individual cell clonotypes, we found that 78% of clonotypes recovered using nanovials had at least one TCR permutation with antigen-specific reactivity and the calling of epitope information was 100% correct for those reactive TCRs. The few non-reactive TCRs were tested with the other peptides and found to be unreactive. We also performed GLIPH2 analysis on CMV1-reactive clones and found that some reactive TCRs recovered using nanovials but not tetramers (TCR21 and TCR22) overlapped in a cluster with tetramer/nanovial clones with different CDR3 regions (Amino acid-motif: QTG). Other nanovial-only clones also clustered separately with a different repertoire of CDR3s (motif: SSV) or having the same CDR3 sequences (motif: Single) (Fig. 5B).

**Figure 5.**
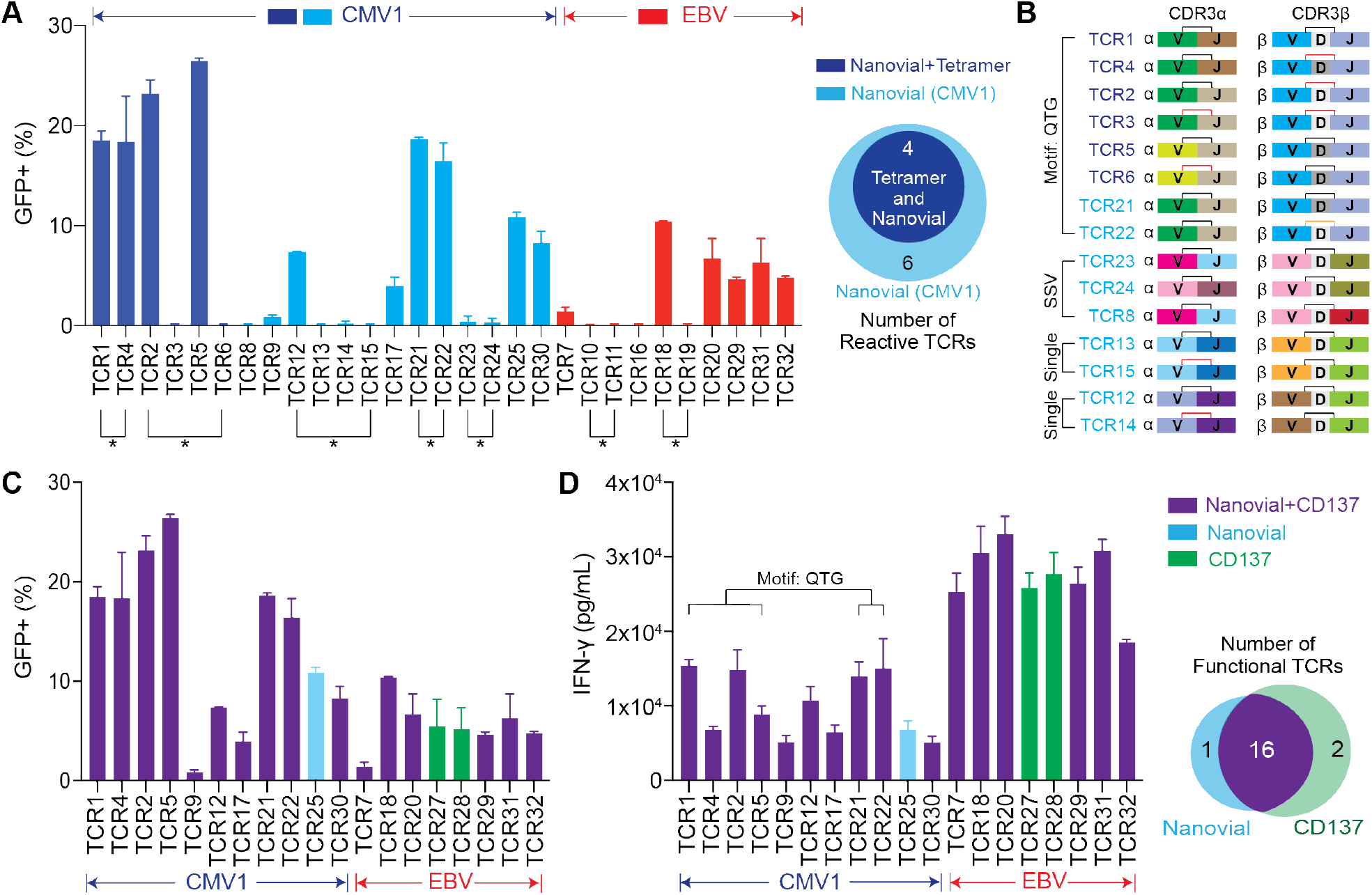
Functional validation of 32 recovered TCRs. (A) The percent of GFP^+^/CD8^+^/murineTCRβ^+^ cells from NFAT-GFP reporter Jurkat cells transduced with respective TCR and exposed to APCs with exogenously added peptides. CMV1-reactive TCRs recovered by both nanovial and tetramer approaches (dark blue) and only nanovials (light blue) or EBV-reactive TCRs recovered by nanovials (red) are plotted. Connecting lines and an asterisk represent TCRs recovered from the same cell clonotype. Venn diagram showing 4 overlapping and 6 additional TCRs recovered by nanovials compared to CMV1 pMHC tetramers. (B) Clustering of recovered TCRs based on GLIPH2 analysis. CDR3s for sorted antigen-specific pools enriched for a subset of motifs (QTG, SSV, Single). Specific V, D, or J sequences are identified with matching colors. Differing CDR3 sequences are indicated with a black or red colored bar, spanning V to J. (C) 19 reactive TCRs are plotted by percent GFP+ cells for TCRs recovered with nanovials and CD137 (purple), and unique TCRs detected by nanovials (light blue) or CD137 (green) alone. (D) IFN-γ secretion measured by ELISA is plotted following exposure of PBMCs transduced with the same 19 reactive TCRs to APCs with added peptide. No secretion was observed from the negative control group, PBMCs transduced with the same vector but without TCRs. TCRs enriched with CDR3s for the same motif are represented by a connecting line and motif nomenclature.

Nanovial and CD137 workflows recovered TCRs with a more extensive range of reactivity (Fig. 5C), including one unique clonotype (TCR25) which was only recovered by nanovials and possessed intermediate reactivity (Fig. 5C). To investigate how combined affinity and functional IFN-γ secretion-based selection on nanovials correlated to secretory function elicited by the recovered TCR sequences in T cells, 19 reactive TCRs identified in the Jurkat-NFAT-GFP assay were transduced into human PBMCs and IFN-γ secretion was measured following exposure to antigen presenting cells (APCs) with exogenously added cognate peptides. We found that all 19 TCRs tested were able to specifically produce secreted IFN-γ (>5000 pg/mL) when stimulated by exogenous peptides (Fig. 5D). Levels of secreted IFN-γ in PBMCs were not directly correlated to GFP activation signals when tested in Jurkat-NFAT-GFP (R^2^=0.10). Nanovial and CD137 approaches were both able to recover TCRs with a range of different potencies, but only nanovials provided matched epitope information.

### Multiplexed secretion-based profiling

T cells engaged with APCs produce multiple cytokines simultaneously to achieve effector functions. We explored the capability of nanovials with additional anti-cytokine capture antibodies to profile multiple cytokines and link this secretion phenotype with surface markers. We first tested on-nanovial assays for detecting multiple cytokines and surface markers simultaneously using primary human T cells activated with PMA and ionomycin (fig. S8A). Both CD8+ and CD4+ cells could be loaded and detected on nanovials with CD8+ cells representing a larger fraction of the population (fig. S8B-C). We sorted populations of cells based on fluorescence peak areas exceeding the positive threshold for each individual cytokine as well as combinations of IFN-γ and TNF-α or IL-2 and IFN-γ (fig. S8D-G). When measuring IFN-γ and TNF-α, the dominant secretion phenotype for CD8 cells was IFΝ-γ (33.5%) and only a small fraction of CD8+ cells secreted TNF-α alone (4.29%) (fig. S8H). About 24% of CD8+ cells were polyfunctional, secreting both IFN-γ and TNF-α simultaneously. On the other hand, CD4+ cells had a larger polyfunctional population (47.5%) and this pattern was consistent when we analyzed for IFN-γ and IL-2 secretion (fig. S8H).

We then sought to test if the multiplexed secretion assay can also be applied to low-potency TCRs targeting tissue antigens since high-affinity T cell clones might get removed during thymic negative selection (*18*). Our previous work led to the discovery of a set of TCRs targeting prostatic acid phosphatase (PAP), a prostate tissue antigen (*19*). Those candidates showed encouraging peptide recognition but had limited cytotoxicity on processed epitopes (*19*). In the context of HLA-A*02:01, TCR128 and 218 transduced PBMCs were loaded onto nanovials conjugated with PAP21 pMHC. TCR156 transduced PBMCs were loaded onto PAP22 pMHC labeled nanovials, and the non-cognate PAP14 pMHC-nanovials as a negative control. Engineered CD3^+^CD8^+^ cells were highly enriched (NGFR^+^% > 90%) for all three tested PAP TCRs when loaded onto nanovials with their cognate pMHC (Fig. 6A, fig. S9A-B), but not when using the non-specific PAP14 pMHC or when loading untransduced cells (fig. S9C-D). CD3^+^CD8^+^ cells representing a variety of secretion phenotypes, including an IFN-γ and TNF-α polyfunctional population, were successfully analyzed and sorted (Fig. 6B-C). We were able to sort and recover the polyfunctional populations with high viability, enabling downstream de novo discovery of TCR sequences based on this unique phenotype in the future.

**Figure 6.**
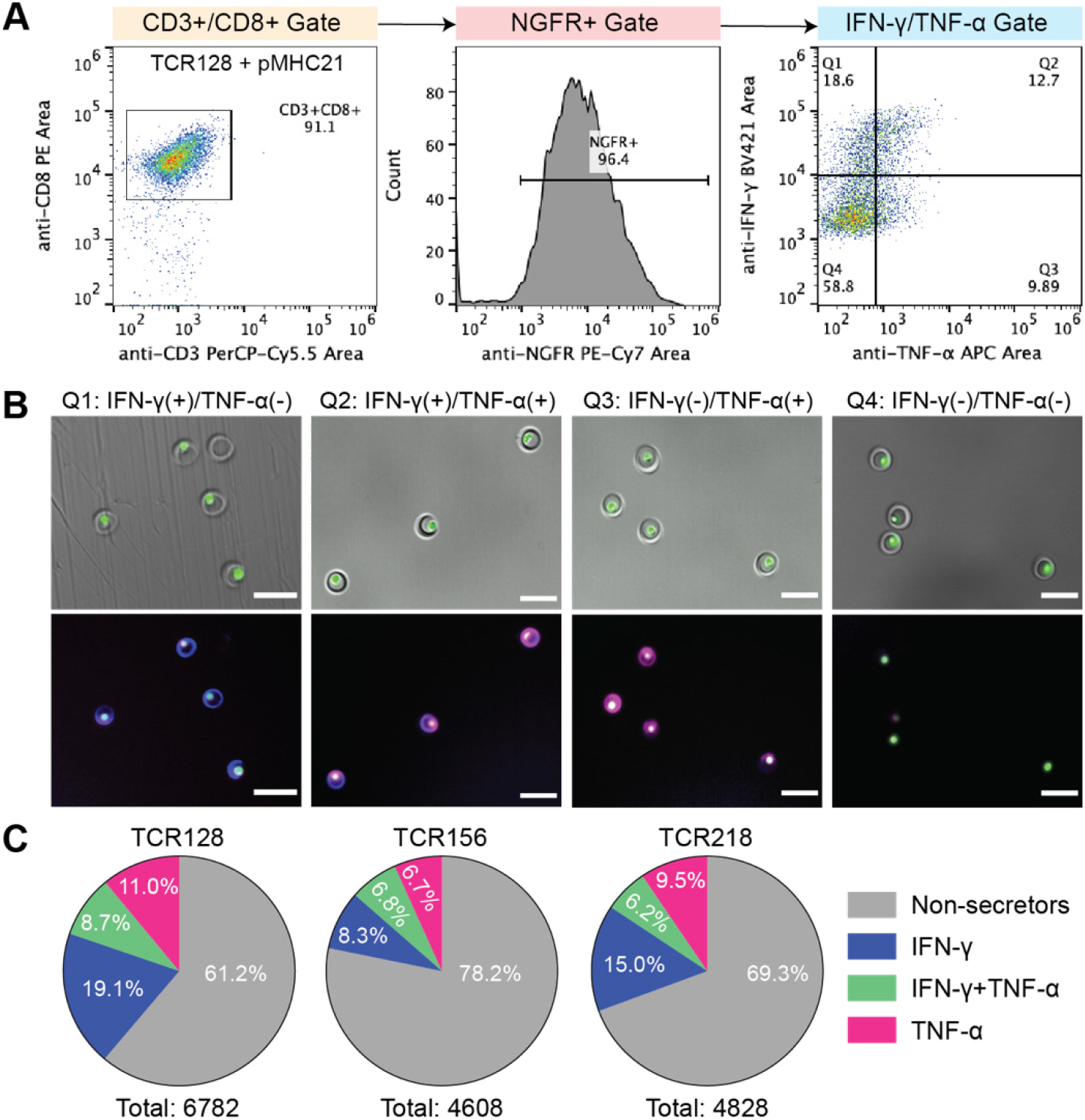
Multiplexed secretion-based profiling of prostate tissue antigen-specific T cells. (A) FACS analysis and sorting gates for identifying functional antigen-specific T cells transduced with TCR128 loaded on HLA-A*02:01 restricted PAP21 pMHC labeled nanovials. IFN-γ and TNF-α secretion signals were analyzed from the CD3^+^/CD8^+^/NGFR^+^ cells. (B) Images of T cells on nanovials that were sorted reflecting each of the four quadrant gates shown in A including an IFN-γ and TNF-α polyfunctional population (Q2). Scale bars represent 50 μm. (C) The population distribution based on secretion phenotype is shown as a pie chart for TCR 128, 218, 156 transduced cells. CD3+CD8+ cells with secretion signal below the background threshold were considered as non-secretors.

## Discussion

Nanovials provide a new tool to sort live antigen-specific T cells based on a combination of TCR affinity and functional response (cytokine secretion) followed by recovery of reactive TCRs and epitope-specific annotation. This approach brings a number of advantages over conventional single-cell cognate T cell isolation platforms. First, nanovials can present pMHC at very high density, providing an initial high avidity enrichment step from a large pool of cells (∼20 million cells in our experiments). Even cells with low affinity TCRs (5G6 and 9D2), which are not easily detectable using tetramer and dextramer staining, were recovered with higher efficiency. The ability to enrich over a broader range of affinity is reflected in the much broader range of CMV-1 reactive TCRs discovered using the nanovial approach compared to tetramer staining. Nanovials were able to recover some previously reported CMV1- and EBV-specific TCR sequences (colored in fig. S10) along with a diverse set of new TCR sequences that were validated to be functional (uncolored) (*20, 21*). This broader range does not come with the trade-off of low purity. High purity screening is supported by both the spiking data and the 78% functional hit rate of TCR clonotypes following re-expression with a matching epitope recall rate of 100%. Other large-scaled pooled barcoded multimer approaches demonstrated a functional hit rate of ∼50% or only assessed functionality of a few TCRs recovered instead of the entire set with 80% accuracy for calling matching epitopes (*22, 23*). Another study showed the ability to link TCR sequences with specific pMHC molecules using barcoded tetramers using single-cell sequencing in a multi-well plate format, although no data was presented on the fraction of the recovered TCR sequences that were reactive when re-expressed (*24*).

The accessibility and compatibility of nanovials with standard FACS and single-cell sequencing instrumentation can accelerate the development of personalized TCR immunotherapies. Epitopes for each recovered TCR are annotated through barcoding, while still being able to recover TCRs over a range of reactivity. The number of pMHCs that can be multiplexed with nanovials is extensible to >40 based on commercial oligonucleotide-barcoding reagents, or ∼1000 using specialized manufacturing approaches (*25*). Since TCR-pMHC interaction is heavily dependent on HLA-subtype restriction, the ability of nanovials to provide TCRs along with matching HLA-restricted epitopes leverages current technology limitations to simultaneously profile a large library of antigen-specific T cells, especially in disease models identified with diverse HLA genotypes, like type 1 diabetes or COVID-19 (*23, 26, 27*).

By screening for TCRs based on the ability of cells to secrete a panel of cytokines, we can further explore links between TCR structure and cellular function and discover therapeutically important TCRs that, for example, are used by different cell subsets, such as regulatory T cells to prevent autoimmune conditions. Recent work has emphasized the importance of functional characterization of TCRs, such as through assaying Ca^2+^ flux upon mechanical engagement of TCRs with pMHC-coated hydrogel beads, a platform that could be synergistic with nanovials to more fully functionally screen TCRs (*28*). Separately, we have shown that oligonucleotide-barcoded antibodies can be used as labels in the nanovial assay format (see secretion-encoded single cell (SEC)-seq workflows) (*29*). Using barcoded antibodies to label secreted cytokines would allow encoding of this cellular function into the single-cell sequencing data set and ranking of TCR sequences based on the amount of cytokine(s) secreted. These types of multiomic studies can ultimately uncover relationships between TCR structure and function for improved efficacy in T cell therapies. Beyond TCRs, the nanovial assay format should be applicable to other screening processes, e.g. for CAR-T cells, CAR-NK cells, TCR-mimics, or bispecific T cell engagers (BiTEs), with minor adjustments, opening up a new frontier in functional screening for cell therapy discovery and development.

## Materials and Methods

### Nanovial fabrication

Polyethylene glycol biotinylated nanovials with 35 μm diameters were fabricated using a three-inlet flow-focusing microfluidic droplet generator, sterilized and stored at 4°C in Washing Buffer consisting of Dulbecco’s Phosphate Buffered Saline (Thermo Fisher) with 0.05% Pluronic F-127 (Sigma), 1% 1X antibiotic-antimycotic (Thermo Fisher), and 0.5% bovine serum albumin (Sigma) as previously reported (*30*).

### Nanovial Functionalization

#### Streptavidin conjugation to the biotinylated cavity of nanovials

Sterile nanovials were diluted in Washing Buffer five times the volume of the nanovials (i.e. 100 μL of nanovial volume was resuspended in 400 μL of Washing Buffer). A diluted nanovial suspension was incubated with equal volume of 200 μg/mL of streptavidin (Thermo Fisher) for 30 minutes at room temperature on a tube rotator. Excess streptavidin was washed out three times by pelleting nanovials at 2000xg for 30 seconds on a Galaxy MiniStar centrifuge (VWR), removing supernatant and adding 1 mL of fresh Washing Buffer.

#### Anti-CD45 and cytokine capture antibody labeled nanovials

Streptavidin-coated nanovials were reconstituted at a five time dilution in Washing Buffer containing 140 nM (20 μg/mL) of each biotinylated antibody or cocktail of antibodies: anti-CD45 (Biolegend, 368534) and anti-IFN-γ (R&D Systems, BAF285), anti-TNF-α (R&D Systems, BAF210), anti-IL-2 (BD Sciences, 555040). Nanovials were incubated with antibodies for 30 minutes at room temperature on a rotator and washed three times as described above. Nanovials were resuspended at a five times dilution in Washing Buffer or culture medium prior to each experiment.

#### pMHC labeled nanovials

MHC monomers with peptides of interest were synthesized and prepared according to a published protocol (*31*). Streptavidin-coated nanovials were reconstituted at a five times dilution in Washing Buffer containing 20 μg/mL biotinylated pMHC and 140 nM of anti-IFN-γ antibody unless stated otherwise. For oligonucleotide barcoded nanovials, 1 μL of 0.5 mg/mL totalseq-C streptavidin (Biolegend, 405271, 405273, 405275) per 6 μL nanovial volume was additionally added during the streptavidin conjugation step.

### Cell culture

#### Human primary T cells

Human primary T cells were cultured as previously reported (*30*).

#### Human donor PBMCs

To prime naïve T cells with peptides of interest, PBMCs from commercial vendors (AllCells) were cultured and processed as previously described with chemically synthesized peptides (>80% purity, Elim Biopharm) (*19*).

#### K562 and Jurkat-NFAT-ZsGreen

K562 (ATCC) and Jurkat-NFAT-ZsGreen (gift from D. Baltimore at Caltech) were cultured in RPMI 1640 (Thermo Fisher) with 10% FBS (Omega Scientific) and Glutamine (Fisher Scientific). 293T (ATCC) was cultured in DMEM (Thermo Fisher) with 10% FBS and Glutamine.

### Nanovial secretion assay general procedure

#### Cell loading onto nanovials

Each well of a 24-well plate was filled with 1 mL of media and 30 μL of reconstituted functionalized nanovials (6 μL of nanovial volume=187,000 total nanovials) was added in each well using a standard micropipette. Cells were seeded in each well and extra culture medium was added to make a total volume of 1.5 mL. Each well was mixed by simply pipetting 5 times with a 1000 μL pipette set to 1000 μL. The well plate was transferred to an incubator to allow cell binding; the volume in each well was pipetted up and down again 5 times with a 200 μL pipette set to 200 μL at 30-minute intervals. After one hour, nanovials were strained using a 20 μm cell strainer to remove any unbound cells and recovered. During this step, any unbound cells were washed through the strainer and only the nanovials (with or without cells loaded) were recovered into a 12-well plate with 2 mL of media by inverting the strainer and flushing with media.

#### Activation, secretion accumulation and secondary antibody staining on nanovials

After cell loading, cells on nanovials in a 12-well plate were activated via 10 ng/mL PMA (Sigma) and 2.5 μM ionomycin (Sigma) or the pMHCs on the nanovials for three hours in the incubator. Each sample was recovered in a conical tube with 5 mL wash buffer and centrifuged for 5 minutes at 200xg. Supernatant was removed and nanovials were reconstituted at a ten-fold dilution in Washing Buffer containing detection antibodies to label secreted cytokines and/or cell surface markers. Concentrations of fluorescent antibodies per 187,000 nanovials (∼ 6 μL nanovial volume) are listed in Table 1, unless otherwise stated. A typical experiment used 30 μL of nanovials, which were incubated with 5X the volumes of antibodies listed in Table 1 (i.e. 25 μL anti-IFN-γ BV421, 10 μL anti-CD3 PerCP Cy5.5 and 10 μL anti-CD8 PE) at the total reaction volume of 300 μL. Nanovials were incubated with the detection antibody cocktail at 37°C for 30 minutes, protected from light. After washing nanovials with 5 mL of Washing Buffer, nanovials were resuspended at a 50-fold dilution in Washing Buffer and transferred to a flow tube.

**Table 1.** Secondary antibody concentrations per 6 μL nanovial volume.

| Calcein AM | Anti-IFN-γ<br>BV421 | Anti-TNF-α<br>APC | Anti-IL-2 APC | Anti-NGFR<br>PE-Cy7 | Anti-CD8<br>PE | Anti-CD3<br>PerCP Cy5.5 | Anti-CD3<br>APC Cy7 |
| --- | --- | --- | --- | --- | --- | --- | --- |
| Thermo Fisher<br>C3099 | Biolegend<br>502532 | Biolegend<br>502912 | BD Sciences<br>554567 | Biolegend<br>345110 | Invitrogen<br>1208842 | Biolegend<br>300430 | Biolegend<br>300426 |
| 0.3 μM | 5 μL of<br>100 μg/mL | 5 μL of<br>100 μg/mL | 5 μL of<br>200 μg/mL | 2 μL of<br>100 μg/mL | 2 μL of<br>125 μg/mL | 2 μL of<br>100 μg/mL | 2 μL of<br>200 μg/mL |

#### Flow cytometer analysis and sorting

All flow cytometry analysis and sorting were performed using the SONY SH800 cell sorter equipped with a 130 micron sorting chip (SONY Biotechnology). The cytometer was configured with violet (405 nm), blue (488 nm), green (561 nm) and red (640 nm) lasers with 450/50 nm, 525/50 nm, 600/60 nm, 665/30 nm, 720/60 nm and 785/60 nm filters. Standard gain settings for different sensors are indicated in table 2 below and gains were adjusted depending on the fluorophores used. In each analysis, samples were compensated using negative (blank nanovials) and positive controls (1000 ng/mL recombinant cytokine captured nanovials labeled with each fluorescent detection antibody or cells stained with each surface marker). Nanovial samples were diluted to approximately 623 nanovial/μL in Washing Buffer for analysis and sorting. Drop delay was configured using standard calibration workflows and single-cell sorting mode was used for all sorting as was previously determined to achieve the highest purity and recovery (*32*). A sample pressure of 4 was targeted. The following order of gating strategy was used to identify cells on nanovials with strong secretion signal: 1) nanovial population based on high forward scatter height and side scatter area, 2) calcein AM positive population, 3) cell surface marker positive population (CD3, CD8, CD4 or NGFR), 4) cytokine secretion signal positive population based on fluorescence peak area and height.

**Table 2.** Common gain settings used for analysis and sorting.

| Sensor | FSC | BSC | FL1 | FL2 | FL3 | FL4 | FL5 | FL6 |
| --- | --- | --- | --- | --- | --- | --- | --- | --- |
| Gain | 1 | 26% | 28% | 22% | 28% | 30% | 32% | 32% |

### Dynamic range of cytokine detection on nanovials with a combination of antibodies

Nanovials were labeled with biotinylated antibodies (140 nM anti-CD45 and 140 nM anti-IFN-γ or anti-TNF-α) using the modification steps mentioned above. Each sample of cytokine capture antibody-labeled nanovials was incubated with 0, 10, 100, or 1000 ng/mL of recombinant human IFN-γ (R&D Systems, 285IF100) and TNF-α (R&D Systems, 210TA020) for 2 hours at 37°C. Excess proteins were removed by washing nanovials three times with Washing Buffer. Nanovials were pelleted at the last wash step and incubated with anti-IFN-γ BV421 and anti-TNF-α APC as described in secondary antibody staining procedure and Table 1. Following washing three times, nanovials were reconstituted at a 50 times dilution in the Washing Buffer and transferred to a flow tube. Fluorescent signal on nanovials was analyzed using a cell sorter with sensors and gains mentioned in the flow cytometer analysis and sorting section.

### Maximum binding of pMHC on nanovials

Streptavidin coated nanovials were functionalized with biotinylated HLA-A*02:01 restricted NY-ESO-1 pMHC by incubating at various concentrations (0, 20, 40, 80, 90, 100 μg/mL) and washed three times as described in Nanovial Functionalization section. Nanovials were reconstituted at a ten-fold dilution in Washing Buffer containing 2 μL of 100 μg/mL anti-HLA-A2 FITC antibody (Biolegend, 343304) and incubated for 30 minutes at 37°C. After washing three times with Washing Buffer, mean fluorescence intensity was measured by flow cytometry.

### Single-T cell loading and statistics

Nanovials labeled with anti-CD45 antibodies were prepared using the procedures described above. To test cell concentration dependent loading of nanovials 0.15 × 10^6^ (0.8 cells per nanovial), 0.3 × 10^6^ (1.6 cells per nanovial), and 0.47 × 10^6^ (2.4 cells per nanovial) of cell tracker deep red stained human primary T cells were each seeded onto 187,000 nanovials in a 24-well plate and recovered as described above. Loading efficiency was analyzed using a custom image analysis algorithm in MATLAB. The software measured the total number of nanovials in each image frame, then the number of cells in each nanovial was manually counted to record the total number of nanovials with 0, 1 or 2 or more cells (n > 2000). For comparing loading with different cell binding motifs, nanovials were labeled with 140 nM of each biotinylated antibody: anti-CD3 (Biolegend, 317320), anti-CD3 and anti-CD28 (Biolegend, 302904), or anti-CD45. Nanovials were seeded with 0.3 million cells in each well. To determine the effect of increased anti-CD45 concentration on nanovials, nanovials were labeled by incubating with 0, 70, 140, or 210 nM of anti-CD45 antibodies and seeded with 0.3 million cells in a 24-well plate. After cell binding and recovery of nanovials, the number of cells in each nanovial was analyzed using the same image analysis algorithms mentioned above (n > 2000).

### Analysis and sorting of human primary T cells based on secretion level

Nanovials were sequentially coated with streptavidin as described above and incubated with a solution of biotinylated antibodies (140 nM anti-CD45 and anti-IFN-γ, anti-TNF-α or anti-IL-2). 0.3 million human primary T cells were seeded on nanovials as described above and recovered into a 12-well plate in 2 mL of T cell expansion medium with PMA and ionomycin, followed by 3 hours of activation. Secreted cytokines (IFN-γ, ΤΝF-α, IL-2) were labeled with fluorescent detection antibodies at concentrations described in Table 1 and cells were stained with calcein AM viability dye. After resuspending nanovials at 50-fold dilution in Washing Buffer, a small fraction of sample was transferred to a 96-well plate to be imaged using a fluorescence microscope prior to sorting. Pre-sort images were analyzed by custom image analysis algorithms in MATLAB. Fluorescence intensity profiles were calculated along a line segment manually defined around the cavity of nanovial. The intensity peak height and the area under the intensity profile were then evaluated to find the peak area over height aspect ratio. Samples were analyzed using a cell sorter based on a combination of fluorescence area and height signals. To sort live single cells based on secretion signal, nanovials with calcein AM staining were first gated and high, medium, or low secretors were sorted by thresholding the fluorescence area and height signals. Sorted samples were imaged with a fluorescence microscope to validate the enrichment of nanovials based on the amount of secreted cytokine captured on the nanovials.

### Capture, activation, and expansion of antigen-specific T cells on pMHC-labeled nanovials

To determine the effect of pMHC concentration on antigen-specific T cell capture efficiency, streptavidin-coated nanovials were functionalized by incubation with different concentrations of biotinylated HLA-A*02:01 NY-ESO-1 pMHCs (10, 20, 40, 80 μg/mL) and seeded with 0.3 million 1G4 TCR transduced PBMCs or untransduced PBMCs. After straining and recovery, samples were stained with calcein AM and anti-NGFR PE Cy7 antibody as described above. The fractions of nanovials with live cells and NGFR positive cells were measured by a cell sorter. To test if activation was specific to the presence of pMHCs on nanovials, 1G4 transduced PBMCs were loaded onto anti-IFN-γ antibody and pMHC or anti-CD45 labeled nanovials. Following 3 hours of activation, nanovial samples were stained with anti-IFN-γ BV421 and anti-NGFR PE Cy7 antibodies. The fraction of nanovials with NGFR positive cells and secretion signal was identified using flow cytometry. Secretion signal from 1G4 PBMCs on pMHC labeled nanovials was measured at 0, 3, 6, and 12 hour time points. For detachment and expansion of antigen-specific T cells post-sort, 1G4 PBMCs loaded onto pMHC nanovials were sorted based on calcein AM and NGFR signal and reconstituted with 0.75 mL media and 0.25 mL of 10 mg/mL Collagenase Type II solution (STEMCELL Technologies), followed by a 2 hour incubation at 37°C. Samples were vortexed 3 times at 20 second intervals and strained through a 20 μm strainer to remove empty nanovials. Cells cultured for 5 days were stained with 0.3 μM calcein AM and 0.02 mg/mL of propidium iodie or fluorescent anti-NGFR antibody and imaged using fluorescence microscopy or analyzed using flow cytometry for NGFR expression.

### Enrichment of antigen-specific T cells on pMHC-labeled nanovials

1G4 transduced PBMCs were diluted with untransduced PBMCs at different dilution ratios. At 1:30, 1:300 and 1:3000 dilution, a total of 5 million cells were seeded with 0.4 million HLA-A*02:01 NY-ESO-1 pMHC labeled nanovials while a total of 20 million cells were seeded with 1.5 million nanovials for 1:30000 dilution. As a negative control, 5 million and 20 million untransduced cells were seeded with 0.4 and 1.5 million pMHC-labeled nanovials respectively. Following recovery and 3 hours of activation, samples were stained with calcein AM and a cocktail of detection antibodies: anti-CD3 PerCp Cy5.5, anti-CD8 PE, anti-IFN-γ BV421 and anti-NGFR PE Cy7 at concentrations described in Table 1. Using a cell sorter, samples were gated and analyzed in the following order: 1) nanovials using forward scatter height and side scatter area, 2) calcein AM, CD3 and CD8 positive cells, 3-1) NGFR positive cells, 3-2) NGFR positive cells with IFN-γ signal. To calculate the enrichment ratio, the number of final NGFR positive cells at each dilution was first subtracted by the number of non-specific NGFR positive cell count observed for untransduced cells and divided by the total calcein AM, CD3, and CD8 positive cells to find a final fraction of NGFR positive cells as shown in Equation 1.1 and 1.2.

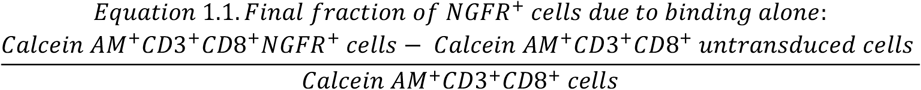

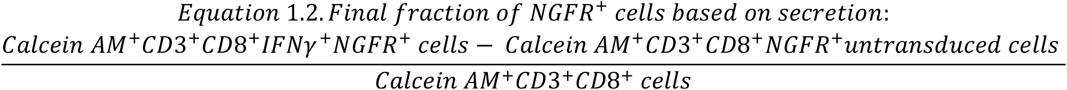

The enrichment ratio at each spiking ratio was then calculated by dividing the final fraction of NGFR positive cells by the initial NGFR positive population using Equation 2 below. Initial fraction of NGFR+ cells was 3.33%, 0.33%, 0.033%, 0.0033% for 1:30, 1:300, 1:3000, and 1:30000, respectively.

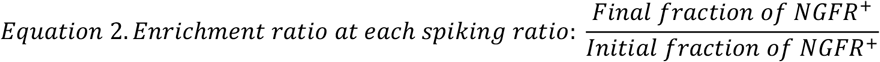

### Enrichment of NY-ESO-1 TCR-transduced cells of various affinities

Each TCR (1G4, 3A1, 4D2, 5G6, 9D2) transduced into PBMCs or untransduced cells were loaded onto HLA-A*02:01 NY-ESO-1 pMHC and anti-IFN-γ labeled nanovials and activated for 3 hours. Recovered samples were stained with a cocktail of detection antibodies (calcein AM, anti-CD3 PerCP Cy5.5, anti-CD8 PE, anti-NGFR PE Cy7, anti-IFN-γ ΒV421) as described above. In parallel, PBMCs transduced with each TCR transduced PBMCs were stained with dual-color commercial HLA-A*02:01 NY-ESO-1 tetramers (MBL International, TB-M105-1 and TB-M105-2), anti-CD3 PerCP Cy5.5, and anti-CD8 PE antibodies as previosly reported (*13*). Using flow cytometric analysis, detection efficiency was calculated as the number of cells bound on nanovials or cells on nanovials with IFN-γ secretion signal divided by the total NGFR positive population in the loaded sample. For tetramer stained samples, detection efficiency was calculated as the number of dual-color tetramer positive population divided by the the total NGFR positive population in the sample. The purity of nanovial sample was calculated as the fraction of NGFR positive population compared to all cells in the calcein AM, CD3 and CD8 positive gate on nanovials or the fraction of the NGFR positive population compared to all cells in the calcein AM, CD3 and CD8 positive gate with secretion signal. The purity of the tetramer stained samples was calculated as the fraction of the NGFR positive population compared to all cells in the CD3 and CD8 positive gate with dual-color tetramer signal. To determine the enrichment ratio, PBMCs transduced with each TCR were diluted into untransduced PBMCs at different dilution ratios (1:10, 1:100, 1:1000). A total of 2 million cells were seeded with 0.4 million pMHC-labeled nanovials for each condition. Following recovery and 3 hours of activation, samples were stained and analyzed using flow cytometry to caculate enrichment ratio as described above.

### Isolation of antigen-specific T cells using nanovials, tetramers and CD137 staining

Nanovials were functionalized with HLA-A*02:01 restricted pMHCs targeting cytomegalovirus pp65, cytomegalovirus IE1 or Eptsein-Barr virus BMLF1 with corresponding totalseq-C streptavidin barcodes C0971, C0972, C0973 (Biolegend, 405271, 405273, 405275) as described above. All sets of functionalized nanovials were pooled together as one nanovial suspension (a total of 0.75 million nanovials). PBMCs were activated for 7 days with peptides associated with each antigen (CMV1: pp65/NLVPMVATV, CMV2: IE1/VLEETSVML, EBV: BMLF1/GLCTLVAML) as reported previously (*19*). 5 million activated PBMCs were loaded onto the pooled nanovial suspension. Following recovery and activation on nanovials for 3 hours, samples were stained with viability dye and a cocktail of detection antibodies (calcein AM, anti-CD3 APC Cy7, anti-CD8, anti-IFN-γ). Using a cell sorter, viable CD3 and CD8 cells on nanovials with IFN-γ secretion signal were sorted. In parallel, 5 million activated PBMCs were each stained with a surface activation marker (CD137) or CMV1 pMHC tetramers and sorted as previously reported (*13*). All sorted samples were reconstituted in 18 μL of 1X PBS containing 0.04% BSA.

### Recovery of TCRs using single-cell TCR αβ sequencing

The standard protocol for 10X Chromium single cell 5’ and V(D)J enrichment with feature barcodes was followed unless otherwise noted. Sorted samples reconstituted at 18 μL were loaded into the 10X Chromium Next GEM Chip K for partitioning each nanovial or cell into droplets containing primers specific for the constant region of the V(D)J locus allowing the PCR amplification and enrichment of matched α and β TCR sequences for individual cell barcoded cDNA. Single-cell TCR V(D)J and feature barcode libraries were constructed using the manufacturer-recommended protocol by the UCLA Technology Center for Genomics & Bioinformatics. Libraries were then sequenced on NextSeq 500 Mid Output with 2×150 bp (Illumina). The Cell Ranger VDJ pipeline was used for sample de-multiplexing and barcode processing.

### Clustering of recovered TCRs using GLIPH2 Analysis

CDR3αβ, Vβ, Jβ and frequency of sequences were submitted for analysis on GLIPH2 online tool, following their standard protocol (http://50.255.35.37:8080/) (*33*). HLA-A*02:01 was inputted as HLA input file. Output clustering data was exported and clusters with same motif were represented in Figure 5A.

### Functional validation of recovered TCR sequences

To measure antigen-specific reactivity of recovered TCR sequences, TCRs were expressed and screened in Jurkat-NFAT-GFP cells as described previously (*34*). Paired TCR alpha and beta chains of interest were cloned into a retroviral pMSGV construct as previously described (*17*). PBMCs for retroviral transduction were processed and cultured according to our rececent publication(*19, 34*). To assess function of the transduced TCRs in human PBMCs, TCR expressing cells were mixed with K562-A2 cells at a ratio of 1:2 (Effector:Target) in the RPMI media and supplemented with 1 µg/ml of anti-CD28/CD49d antibodies (BD Biosciences, 347690) and 1 µg/ml of cognate peptides. For PBMCs, supernatants were collected after 48 hours and analyzed by ELISA (BD Biosciences) to estimate IFN-γ concentration. PBMCs transduced by the vector without a TCR was used as a negative control.

### Multiplexed secretion based profiling to identify polyfunctional T cells

#### Linking cell surface markers to secretion phenotype

Streptavidin-coated nanovials were decorated with biotinylated antibodies (140 nM of anti-CD45, anti-IFN-γ and anti-TNF-α or anti-CD45, anti-IFN-γ and 140 nM anti-IL-2). Negative control nanovials were prepared by labeling nanovials only with anti-CD45 antibody without any cytokine capture antibodies. 0.5 million human primary T cells were loaded onto nanovials and recovered in T cell expansion medium containing 10 ng/mL PMA and 500 ng/mL ionomycin. Following 3 hours of activation, secreted cytokines were stained with fluorescent detection antibodies (anti-IFN-γ BV421, anti-TNF-α APC, anti-IL-2 APC) and cells were stained with 0.3 μM calcein AM, 5 μL of 25 μg/mL anti-CD4 PE (Biolegend, 344606) and 5 μL of 100 μg/mL anti-CD8 Alexa Fluor 488 (Biolegend, 344716) per 6 μL nanovial volume. Using a cell sorter, CD4 or CD8 cells on nanovials with secretion signal were evaluated by first creating quadrant gates based on the negative control sample (nanovials only labeled with anti-CD45 antibody). Q1 was defined as nanovials with only IFN-γ secreting cells. Q2 was nanovials with polyfunctional T cells that secreted both cytokines (IFN-γ and TNF-α or IL-2). Q3 was nanovials with either TNF-α or IL-2 secreting cells while Q4 was nanovials with non-secretors. Nanovials in each quadrant were sorted and imaged with a fluorescence microscope to quantify enrichment of each cell type and their associated secretion characteristics.

#### Multiplexed secretion-based profiling of cancer-specific cognate T cell

PBMCs were transduced with prostate acid phosphatase specific TCRs (TCR128, 156, 218) as previously described (*19*). Streptavidin-coated nanovials were functionalized with biotinylated anti-IFN-γ, anti-TNF-α and pMHC targeting each TCR: PAP21 for both TCR128 and TCR218, and PAP22 for TCR156. As a negative control, non-cognate pMHC (PAP14 for TCR156) labeled nanovials were also prepared. PBMCs transduced with each TCR were separately loaded onto nanovials and activated for 3 hours, followed by secondary staining with anti-CD3 PerCp Cy5.5, anti-CD8 PE, anti-IFN-γ ΒV421, anti-TNF-α APC, anti-NGFR PE Cy7 and calcein AM. Using a cell sorter, CD3, CD8 and NGFR positive cells on nanovials with each secretion phenotype (IFN-γ, TNF-α, polyfunctional, non-secretors) were sorted and imaged using a fluorescence microscope.

## Acknowledgements

We would like to thank other members of Di Carlo lab and Witte lab for helpful comments and discussion in preparation of this manuscript. We thank UCLA Jonsson Comprehensive Cancer Center (JCCC) and Center for AIDS Research Flow Cytometry Core Facility. We thank UCLA Technology Center for Genomics & Bioinformatics for performing sequencing services. We thank Jamie Spangler and Monika Kizerwetter for helpful discussions.

## Funding

National Institutes of Health (5R21CA256084-02)

Parker Institute for Cancer Immunotherapy (20163828)

California NanoSystems Institute (CNSI) Noble Family Innovation Fund

CNSI Mann Family Foundation Technology Development Fund

## Author contributions

Conceptualization: DK, ZM, DD, ONW

Methodology: DK, ZM, RD, JM, DD, ONW

Investigation: DK, ZM, RD, NT, MN, JM, WT, SL, DC, GB

Formal Analysis: DK, ZM, RD, MN, JM, WT

Visualization: DK, ZM

Funding acquisition: DD, ONW

Writing – original draft: DK, ZM, DD, ONW

Writing – review & editing: DK, ZM, JD, DD, ONW

## Competing interests

J.D. is an employee of Partillion Bioscience which is commercializing nanovial technology. Some of authors are inventors on a patent application assigned to the University of California. J.D., D.D. and the University of California have financial interests in Partillion Bioscience. O.N.W. currently has consulting, equity, and/or board relationships with Trethera Corporation, Kronos Biosciences, Sofie Biosciences, Breakthrough Properties, Vida Ventures, Nammi Therapeutics, Two River, Iconovir, Appia BioSciences, Neogene Therapeutics, 76Bio, and Allogene Therapeutics.

## List of Supplementary Materials

Materials and Methods

Figs. S1 to S10

Supplementary Note

