## Supplementary Information for "Defining T cell receptor repertoires using nanovial-based affinity and functional screening"

### Supplementary Materials for: Defining T cell receptor repertoires using nanovial-based affinity and functional screening

**Authors:** Doyeon Koo[1†], Zhiyuan Mao[2†], Robert Dimatteo[3], Natalie Tsubamoto[1], Miyako Noguchi[4], Jami McLaughlin[4], Wendy Tran[4], Sohyung Lee[3], Donghui Cheng[5], Joseph de Rutte[1,6], Giselle Burton Sojo[4], Owen N. Witte[2,4,5,7,8,9]*, Dino Di Carlo[1,6,10,11]*

**Affiliations:**
[1]Department of Bioengineering, University of California, Los Angeles; Los Angeles, CA 90095, USA.
[2]Department of Molecular and Medical Pharmacology, David Geffen School of Medicine, University of California, Los Angeles; Los Angeles, CA 90095, USA.
[3]Department of Chemical and Biomolecular Engineering, University of California, Los Angeles; Los Angeles, CA 90095, USA.
[4]Department of Microbiology, Immunology and Molecular Genetics, University of California, Los Angeles; Los Angeles, CA 90095, USA.
[5]Eli and Edythe Broad Center of Regenerative Medicine and Stem Cell Research, University of California, Los Angeles; Los Angeles, CA 90095, USA.
[6]Partillion Bioscience; Los Angeles, CA 90095, USA.
[7]Molecular Biology Institute, University of California, Los Angeles; Los Angeles, CA 90095, USA.
[8]Jonsson Comprehensive Cancer Center, University of California, Los Angeles; Los Angeles, CA 90095, USA.
[9]Parker Institute for Cancer Immunotherapy, David Geffen School of Medicine, University of California, Los Angeles; Los Angeles, CA 90095, USA.
[10]Department of Mechanical and Aerospace Engineering, University of California, Los Angeles; Los Angeles, CA 90095, USA.
[11]California NanoSystems Institute; Los Angeles, CA 90095, USA.
†These authors contributed equally to this work.

**Supplementary Materials:**

Figure S1. Nanovial fabrication and functionalization.

Figure S2. Optimization of human primary T cell loading.

Figure S3. Selection of viable T cells based on cytokine secretion.

Figure S4. Optimization of antigen-specific T cell loading, secretion on nanovials and expansion of cells post-isolation.

Figure S5. Flow cytometry sorting gates for identifying functional antigen-specific T cells on nanovials.

Figure S6. FACS analysis and gating strategy for isolation of cognate T cells using nanovials, tetramers or CD137 activation markers.

Figure S7. Determination of the dominant epitope for each clonotype from the number of barcodes detected.

Figure S8. Multiplexed secretion profiling combined with cell surface labeling.

Figure S9. Detailed FACS analysis and gating strategy for multiplexed secretion-based profiling of PAP-specific T cells.

Figure 10. Detailed list of clonotypes recovered from nanovials (frequency >=2) with corresponding V(D)J αβ genes, CDR3 αβ sequences.

Supplementary Note on Area vs. Height Gating for Secretion Detection.

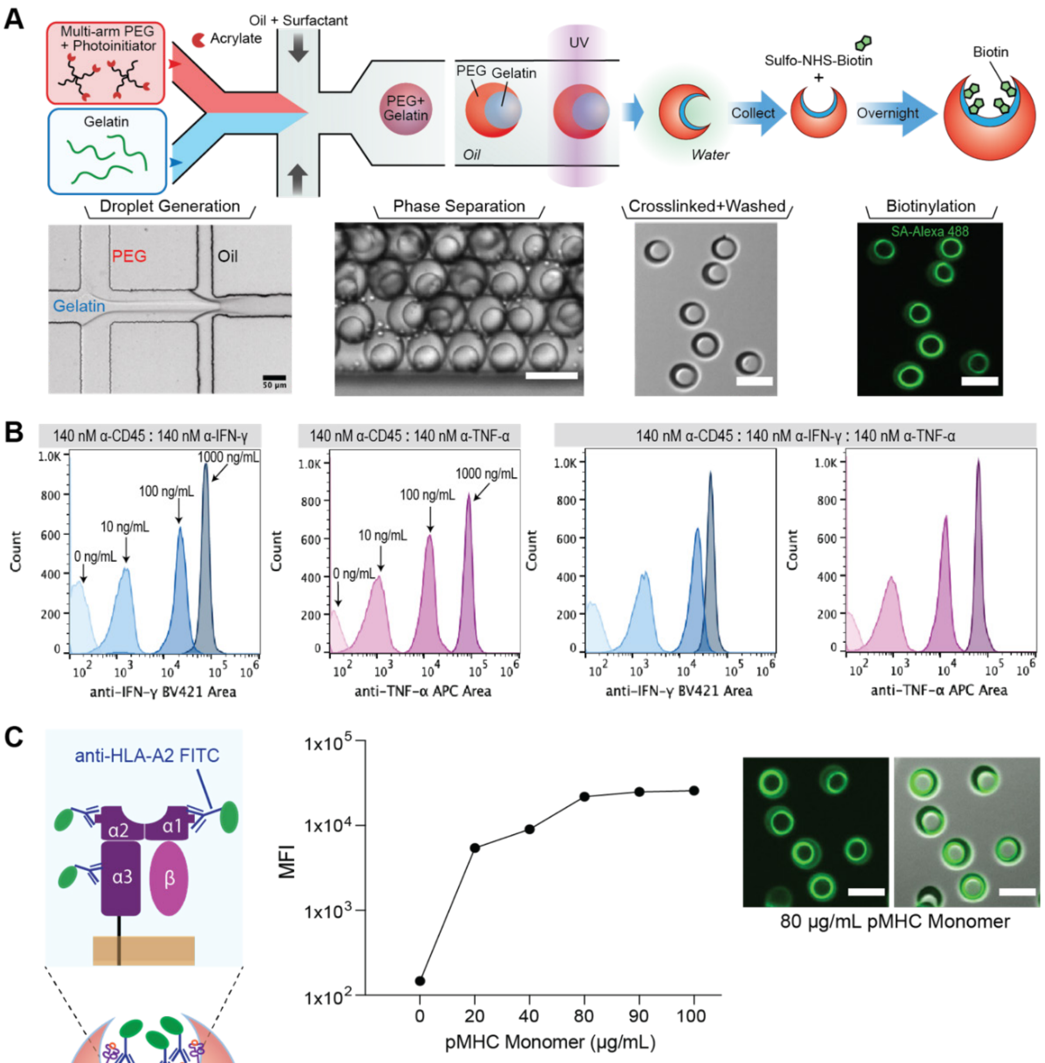


| pMHC Monomer MW | pMHC Monomer Added to Nanovials | Number of Nanovials | pMHC Monomer per Nanovial |
|---|---|---|---|
| 47 kDa | 2.4 µg (80 µg/mL) | 187,000 | 12.8 pg (1.64e8 monomers) |

Figure S1: Nanovial fabrication and functionalization. (A) An aqueous phase comprised of 4-arm-PEG Acrylate and photoinitiator is co-flowed with a gelatin solution in a microfluidic droplet generator. After droplet formation, PEG and gelatin undergo phase separation and are exposed to UV light to cross-link. After collection, nanovials are incubated with sulfo-NHS-biotin to be biotinylated. Localized fluorescence of AlexaFluor488-labeled streptavidin is observed in nanovial cavities. Scale bar represents 50 µm. (B) Flow cytometry fluorescence histograms of nanovials following a dose-dependent cytokine capture assay with different capture antibody combinations. Nanovials were functionalized with anti-CD45 and with one or two cytokine capture antibodies (anti-IFN-γ, anti-TNF-α) and incubated with 0, 10, 100, or 1000 ng/mL of recombinant TNF-α or IFN-γ. The ability of nanovials to detect each individual cytokine was not significantly affected by the presence of other cytokine capture antibodies. (C) Fluorescence intensity based on the concentration of pMHC incubated with nanovials. Nanovials were incubated with 0-100 µg/mL of pMHC monomers and stained with fluorescent anti-HLA-A2 antibodies followed by measurement of fluorescence intensity via SONY SH800 sorter. Signal of loaded pMHC increases linearly up to a concentration of ~80 µg/mL of pMHC monomers.

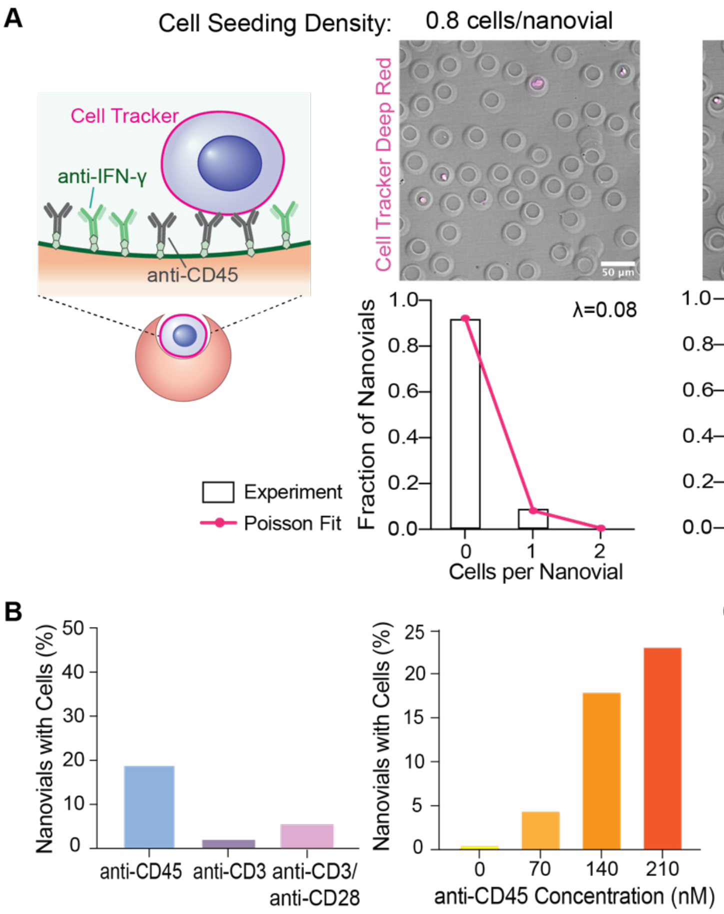


Figure S2: Optimization of human primary T cell loading. (A) Loading of human primary T cells into anti-CD45 and anti-IFN-γ labeled nanovials at different cell number to nanovial ratios. The highest fraction of single-cell loaded nanovials was achieved when cells were seeded at 1.6 cells per nanovial. Increased cell seeding density resulted in a larger fraction of nanovials with two or more T cells. Loading of cells into nanovial cavities followed Poisson statistics. (B) Dependence of cell binding on surface marker target and antibody concentration. Left: Nanovials conjugated with anti-CD45 showed the highest loading efficiency when primary T cells were loaded after initial CD3/CD28 activation in culture. Right: Increased anti-CD45 concentration on nanovials increased cell binding by nearly 6-fold. (C) Effect of cytokine capture antibodies on cell loading. Flow cytometry histograms of cells (calcein AM positive) loaded on nanovials in the presence of anti-IFN-γ or anti-TNF-α antibodies.

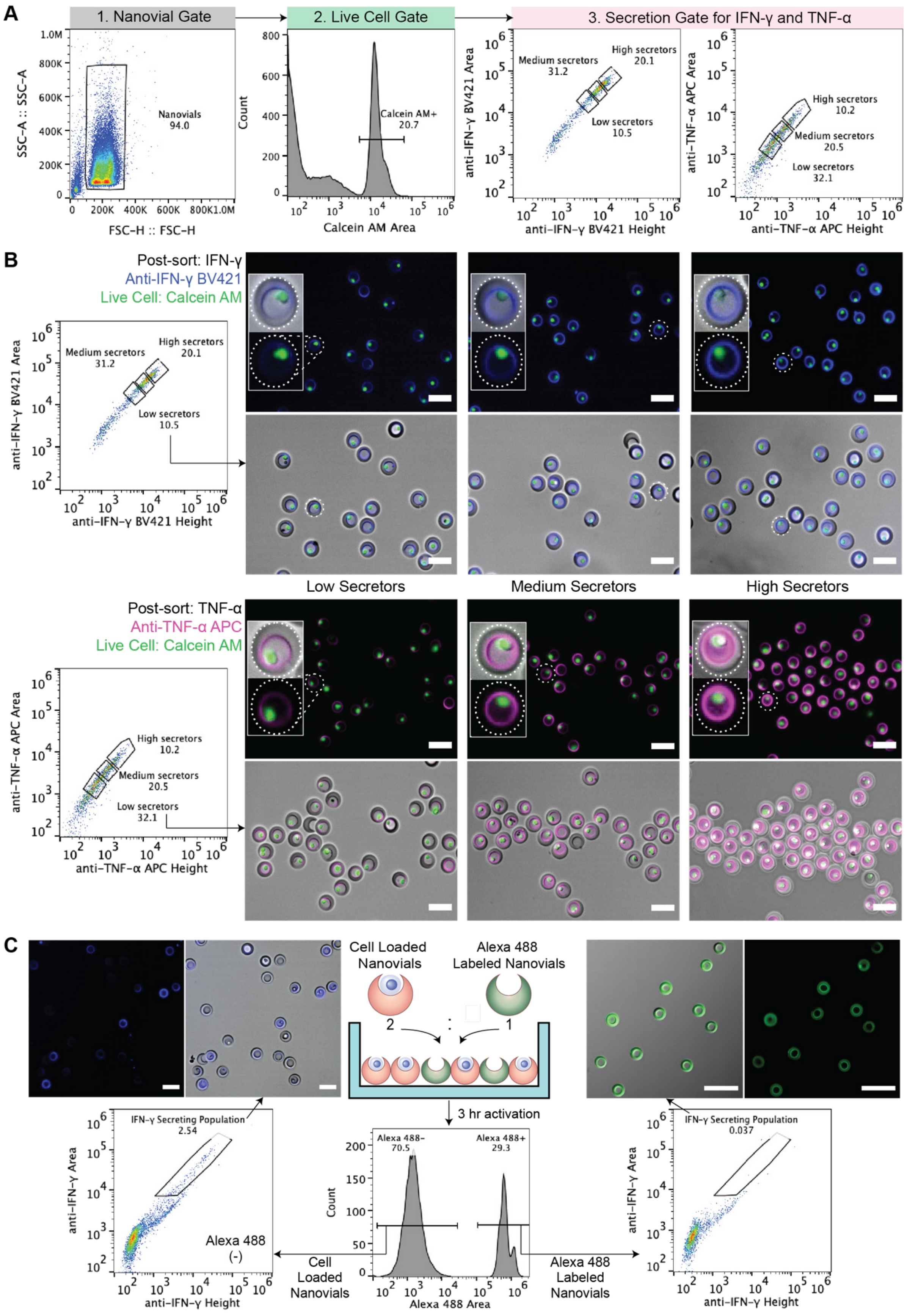
A
1. Nanovial Gate
SSC-A :: SSC-A
FSC-H :: FSC-H
Nanovials 94.0
2. Live Cell Gate
Count
Calcein AM Area
Calcein AM+ 20.7
3. Secretion Gate for IFN-γ and TNF-α
anti-IFN-γ BV421 Area
anti-IFN-γ BV421 Height
Medium secretors 31.2
High secretors 20.1
Low secretors 10.5
anti-TNF-α APC Area
anti-TNF-α APC Height
High secretors 10.2
Medium secretors 20.5
Low secretors 32.1
B
Post-sort: IFN-γ
Anti-IFN-γ BV421
Live Cell: Calcein AM
Low Secretors
Medium Secretors
High Secretors
Post-sort: TNF-α
Anti-TNF-α APC
Live Cell: Calcein AM
C
Cell Loaded Nanovials
Alexa 488 Labeled Nanovials
2
1
3 hr activation
Alexa 488- 70.5
Alexa 488+ 29.3
Count
Alexa 488 Area
IFN-γ Secreting Population 2.54
anti-IFN-γ Area
anti-IFN-γ Height
Alexa 488 (-)
Cell Loaded Nanovials
Alexa 488 Labeled Nanovials
IFN-γ Secreting Population 0.037

Figure S3: Selection of viable T cells based on cytokine secretion. (A) Gates for isolating cells on nanovials with secretion signal. Using a SONY SH800S, the nanovial population was identified in FSC/SSC and further gated for high calcein AM signal. Secretion signal was quantified for this sub-population using the fluorescence peak area vs. peak height (A/H). Single cells were sorted as high, medium, and low secretors based on IFN-γ and TNF-α secretion level. (B) Fluorescence microscopy images of sorted high, medium, and low secretors. Dotted lines in insets outline the nanovial boundaries. Scale bars represent 50 μm. (C) Crosstalk between nanovials. Two nanovial types were introduced together to evaluate crosstalk. Fluorescently labeled nanovials (AlexaFluor 488) without cells were mixed with T cell-loaded nanovials activated with PMA/ionomycin at a ratio of 1:2. Secretion signal was labeled and analyzed on both nanovial types by first gating on the green fluorescence signal on nanovials (Alexa 488 Area). Scatter plots of the cell-loaded nanovial population shows a larger population with high IFN-γ secreting cells compared to nanovials that were not loaded with cells. The percent of cells in the identical gated regions are shown. Scale bars represent 100 μm.

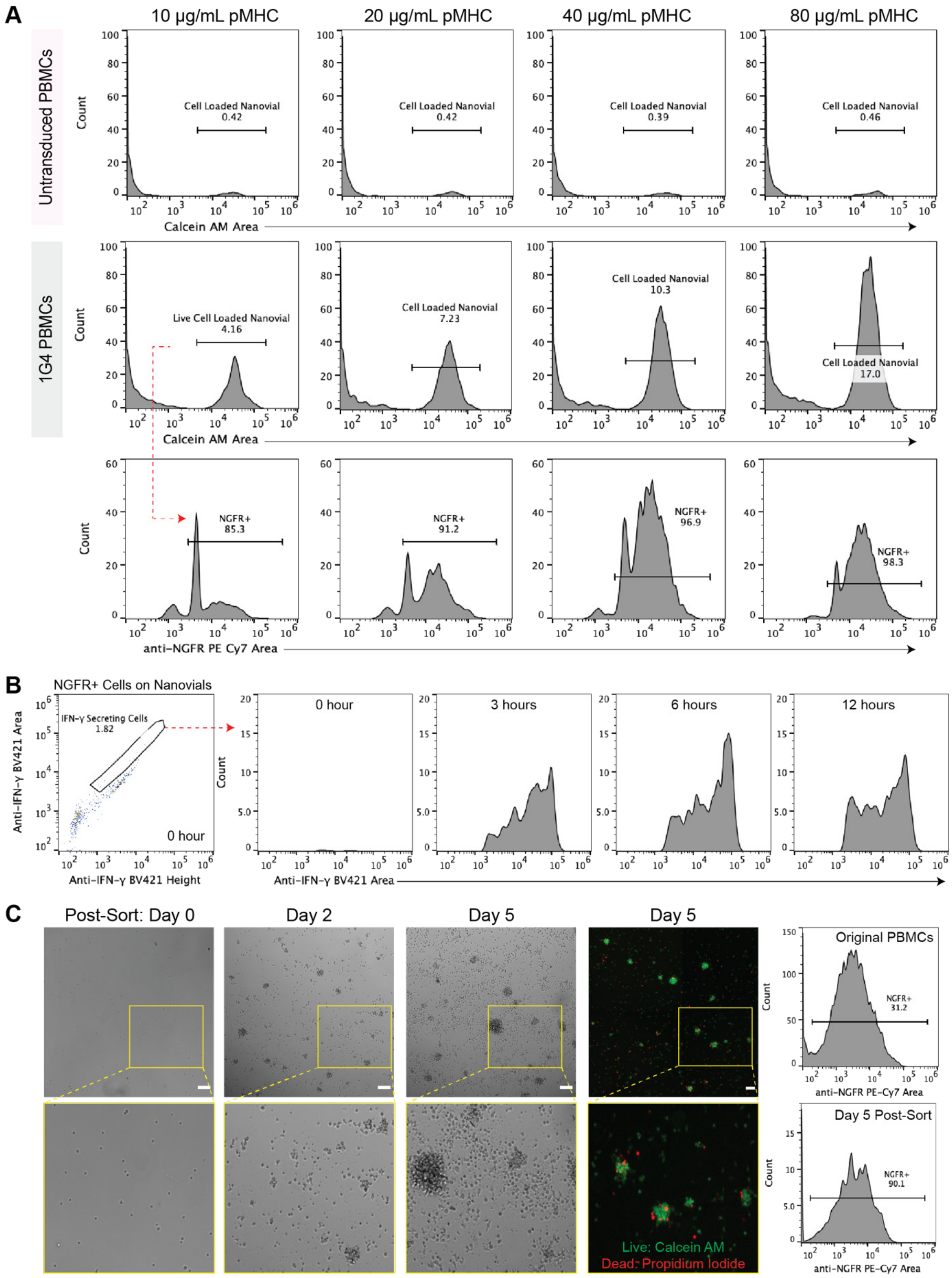


Figure S4: Optimization of antigen-specific T cell loading, secretion on nanovials and expansion of cells post-isolation. (A) Analysis of antigen-specific T cell loading as a function of pMHC concentrations on nanovials. Top flow cytometry histograms show the fraction of untransduced cell loaded nanovials based on calcein AM signal. Bottom shows the fraction of 1G4-transduced cell nanovials and corresponding NGFR positivity of those bound cells (red dashed line). (B) Flow cytometry fluorescence histograms of nanovials following loading of 1G4-transduced PBMCs on pMHC-labeled nanovials. Histograms for IFN-γ secretion are shown as a function of time (0, 3, 6, 12 hours). (C) Images showing expansion of 1G4-transduced T cells following sorting and detachment with collagenase D treatment. Microscopy images show cells expanded in culture over 5 days. (Right) Flow cytometry fluorescence histogram of NGFR levels for the initial (pre-loading onto nanovials) and the expanded population expressing the 1G4 TCR on Day 5. Scale bars represent 50 μm.

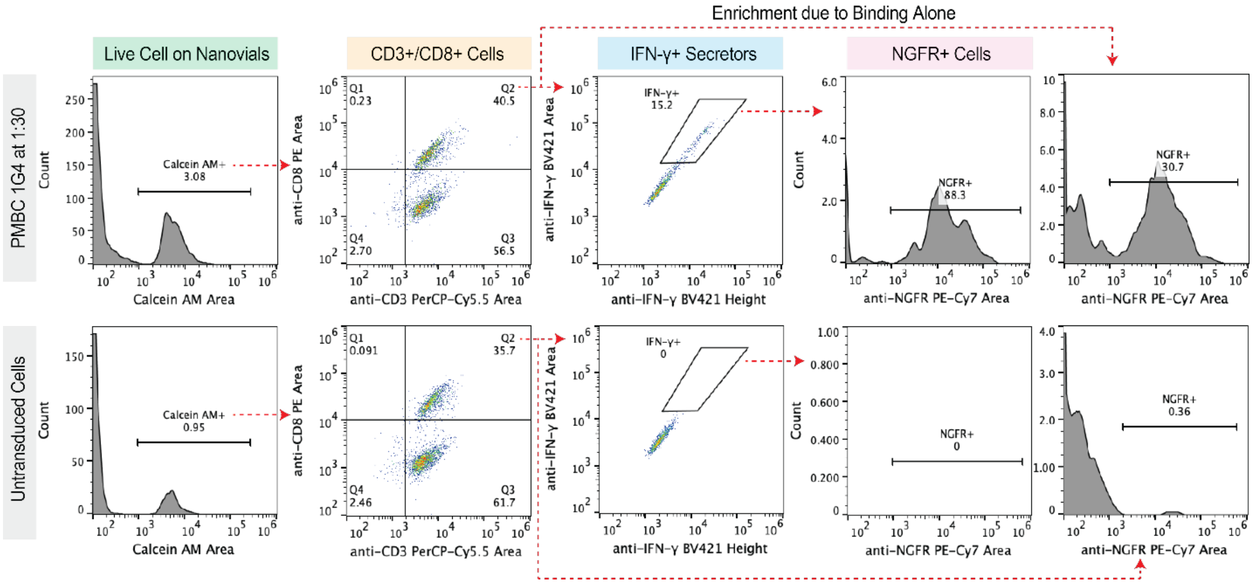


Figure S5: Flow cytometry sorting gates for identifying functional antigen-specific T cells on nanovials. Top plots show 1G4-transduced PBMCs loaded onto NY-ESO-1 pMHC labeled nanovials at a 1:30 dilution with untransduced cells. The bottom plots show untransduced PBMCs on pMHC labeled nanovials. Enriched cells were defined based on gates on viability (calcein AM), CD3 and CD8, and IFN-γ secretion. The NGFR fraction above the background was also determined using a gate on PE-Cy7 area.

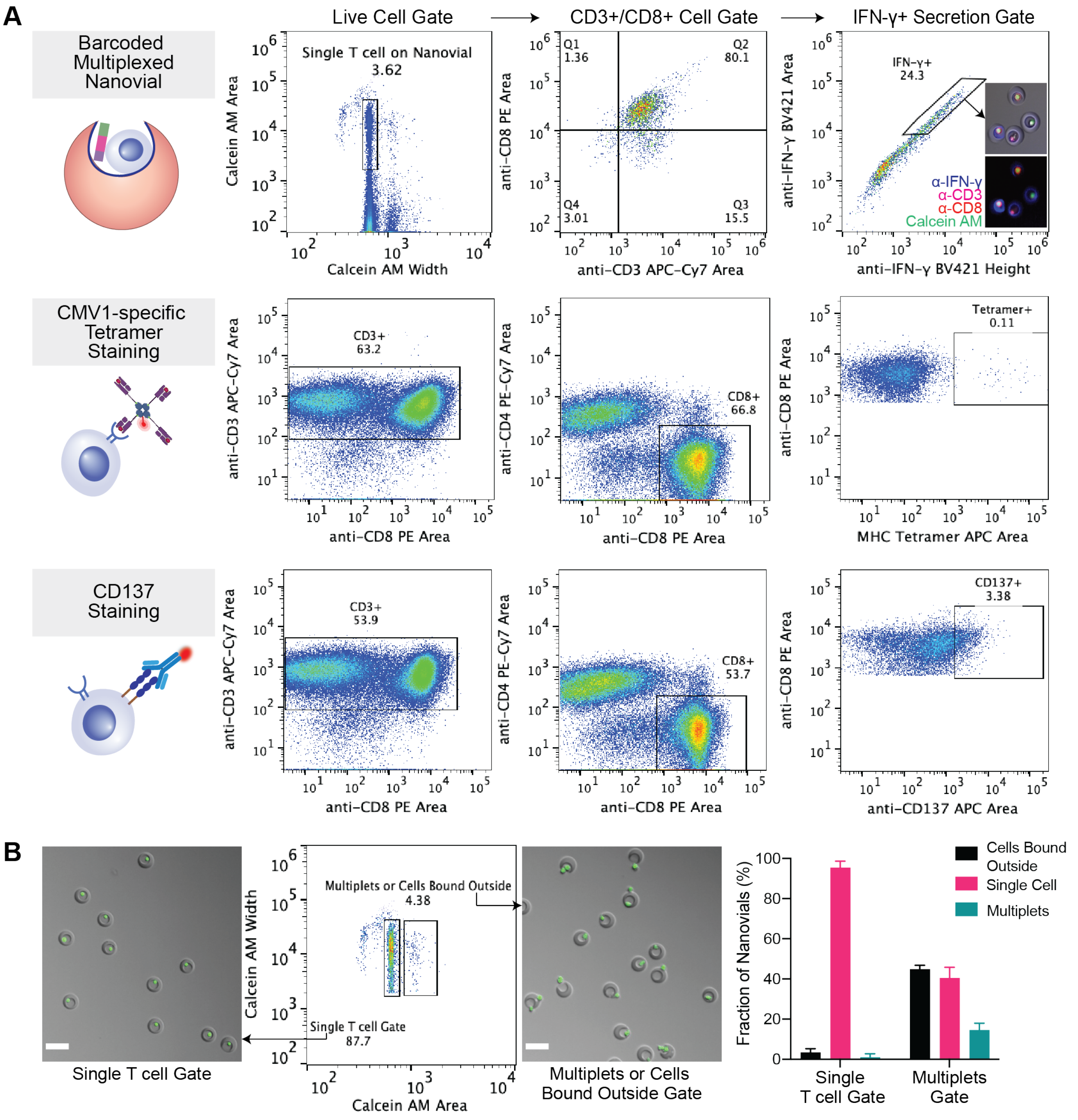


Figure S6: FACS analysis and gating strategy for isolation of cognate T cells using nanovials, tetramers, or CD137 activation markers. (A) Flow scatter plots showing gates for selecting viable CD8+ T cells binding to nanovials and secreting IFN-γ, CD137+ CD8+ T cells, and tetramer+ CD8+ T cells. Microscopy images show representative sorted cells on nanovials. Scale bar represents 50 µm. (B) Scatter plots show calcein AM fluorescence Peak Area vs. Peak Width gates used for isolating single cells on nanovials. Images of sorted events show that the population with a similar peak area but with a shorter peak width contains >90% single cells loaded on nanovials. Data on single-cell occupancy is shown in a bar graph. Scale bars represent 50 µm.

| Clonotype_id | Frequency | Number of Barcodes Detected | Dominant Epitope | Dominant Epitope (%) |
|---|---|---|---|---|
| clonotype0 | 801 | CMV1(597), EBV(10), CMV2(12) | CMV1 | 96.4 |
| clonotype1 | 308 | CMV1(187), EBV(2), CMV2(7) | CMV1 | 95.4 |
| clonotype2 | 83 | CMV1(29), CMV2(2), EBV(1) | CMV1 | 90.6 |
| clonotype3 | 49 | EBV(36), CMV1(3) | EBV | 92.3 |
| clonotype4 | 41 | CMV1(31), EBV(1), CMV2(1) | CMV1 | 93.9 |
| clonotype5 | 35 | CMV1(26) | CMV1 | 100.0 |
| clonotype6 | 26 | EBV(21), CMV1(2) | EBV | 91.3 |
| clonotype11 | 14 | CMV1(13) | CMV1 | 100.0 |
| clonotype8 | 13 | CMV1(3) | CMV1 | 100.0 |
| clonotype9 | 10 | CMV1(6) | CMV1 | 100.0 |
| clonotype7 | 10 | EBV(10) | EBV | 100.0 |
| clonotype13 | 8 | EBV(5) | EBV | 100.0 |
| clonotype12 | 7 | CMV1(5) | CMV1 | 100.0 |
| clonotype10 | 7 | EBV(6) | EBV | 100.0 |
| clonotype14 | 6 | CMV1(5) | CMV1 | 100.0 |
| clonotype20 | 5 | CMV1(3) | CMV1 | 100.0 |
| clonotype22 | 4 | EBV(2) | EBV | 100.0 |
| clonotype18 | 3 | EBV(2) | EBV | 100.0 |
| clonotype26 | 2 | CMV1(1) | CMV1 | 100.0 |
| clonotype51 | 2 | CMV1(2) | CMV1 | 100.0 |
| clonotype45 | 2 | EBV(1) | EBV | 100.0 |
| clonotype35 | 2 | EBV(2) | EBV | 100.0 |
| clonotype36 | 2 | CMV1(1) | CMV1 | 100.0 |
| clonotype34 | 1 | CMV1(1) | CMV1 | 100.0 |
| clonotype48 | 1 | CMV1(1) | CMV1 | 100.0 |
| clonotype69 | 1 | CMV1(1) | CMV1 | 100.0 |
| clonotype77 | 1 | CMV1(1) | CMV1 | 100.0 |
| clonotype78 | 1 | CMV1(1) | CMV1 | 100.0 |
| clonotype80 | 1 | CMV1(1) | CMV1 | 100.0 |
| clonotype93 | 1 | CMV1(1) | CMV1 | 100.0 |
| clonotype106 | 1 | CMV1(1) | CMV1 | 100.0 |
| clonotype129 | 1 | CMV1(1) | CMV1 | 100.0 |
| clonotype145 | 1 | CMV1(1) | CMV1 | 100.0 |
| clonotype165 | 1 | CMV1(1) | CMV1 | 100.0 |
| clonotype168 | 1 | CMV1(1) | CMV1 | 100.0 |
| clonotype179 | 1 | CMV1(1) | CMV1 | 100.0 |
| clonotype229 | 1 | CMV1(1) | CMV1 | 100.0 |
| clonotype241 | 1 | CMV1(1) | CMV1 | 100.0 |
| clonotype16 | 1 | CMV1(2) | CMV1 | 100.0 |
| clonotype19 | 1 | EBV(1) | EBV | 100.0 |
| clonotype21 | 1 | EBV(1) | EBV | 100.0 |
| clonotype50 | 1 | EBV(1) | EBV | 100.0 |
| clonotype58 | 1 | EBV(1) | EBV | 100.0 |
| clonotype99 | 1 | EBV(1) | EBV | 100.0 |
| clonotype134 | 1 | EBV(1) | EBV | 100.0 |
| clonotype148 | 1 | EBV(1) | EBV | 100.0 |
| clonotype177 | 1 | CMV2(1) | CMV2 | 100.0 |
| clonotype192 | 1 | EBV(1) | EBV | 100.0 |
| clonotype213 | 1 | EBV(1) | EBV | 100.0 |

Figure S7: Determination of the dominant epitope for each clonotype from the number of barcodes detected. For clonotypes matched with more than one barcode (clonotype0, 1, 2, 3, 4, 6), the final epitope was determined based on the barcode with the highest fraction, representing >90% of cells represented.

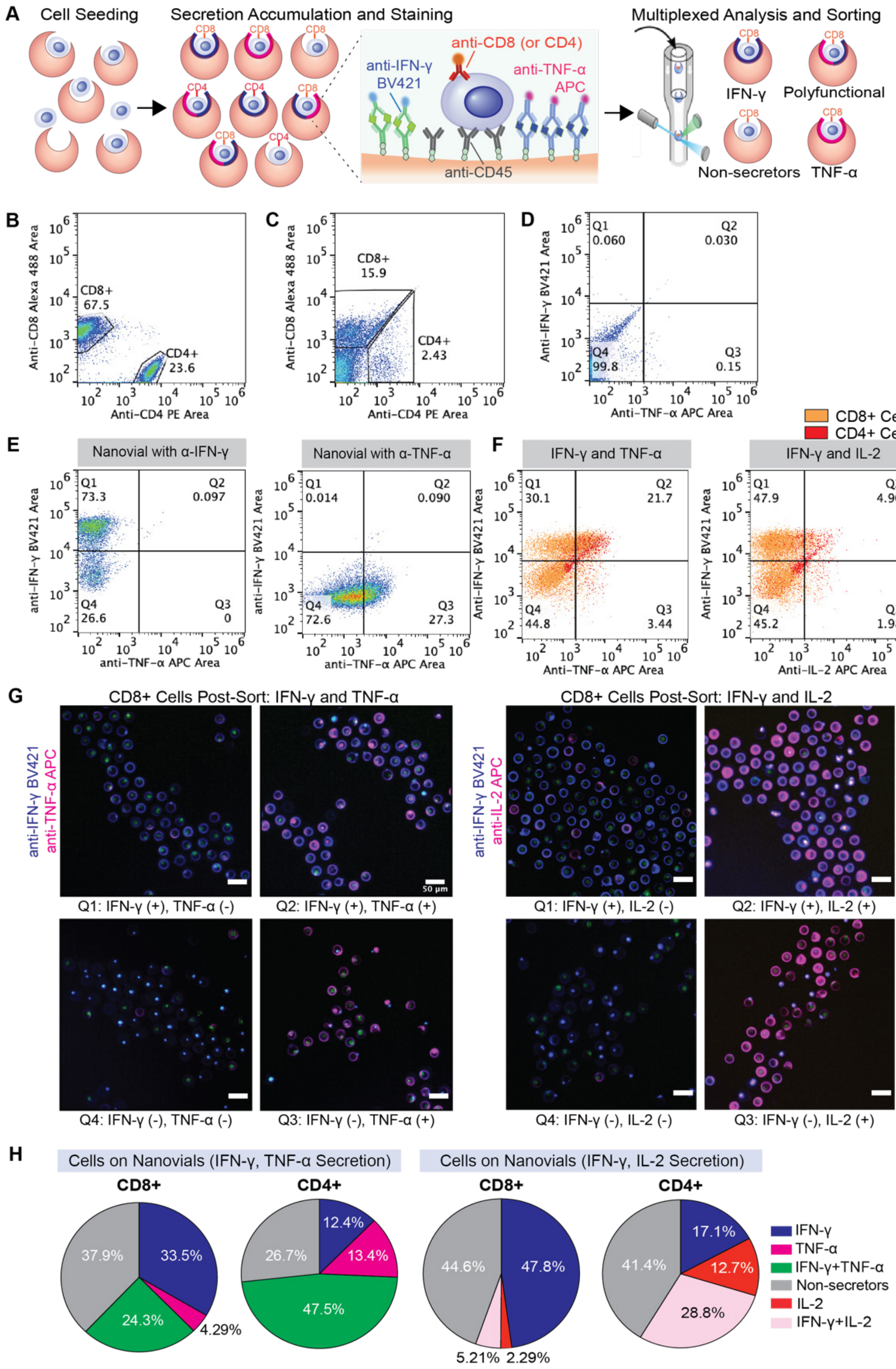


Figure S8: Multiplexed secretion profiling combined with cell surface labeling. (A) Overview of multiplexed profiling of single-T cells based on cytokine secretion and cell phenotype. Human primary T cells loaded on nanovials labeled with two cytokine capture antibodies (anti-IFN-γ and anti-TNF-α, or anti-IFN-γ and anti-IL-2) and anti-CD45 were activated under PMA/ionomycin stimulation. Secreted cytokines and cells were stained with fluorescent detection antibodies, followed by analysis and sorting with a cell sorter. (B) Flow cytometry dot plots showing the distribution of CD8 and CD4 cells in the original cell culture and (C) CD8+ and CD4+ gates for cells on nanovials. (D) Control nanovial dot plots for secretion baseline gates without any secretion capture antibodies and (E) positive threshold gates for IFN-γ and TNF-α on nanovials when using only single cytokine capture antibody. (F) Experimental dot plots showing IFN-γ and TNF-α or IFN-γ and IL-2 secretion from activated CD4+ and CD8+ T cells loaded on nanovials. (G) Single cells were sorted based on CD4+ or CD8+ gates as well as the four quadrant gates shown in F. Scale bars represent 50 μm. (H) A pie chart for CD4+ and CD8+ cells showing the distribution of secretion phenotypes.

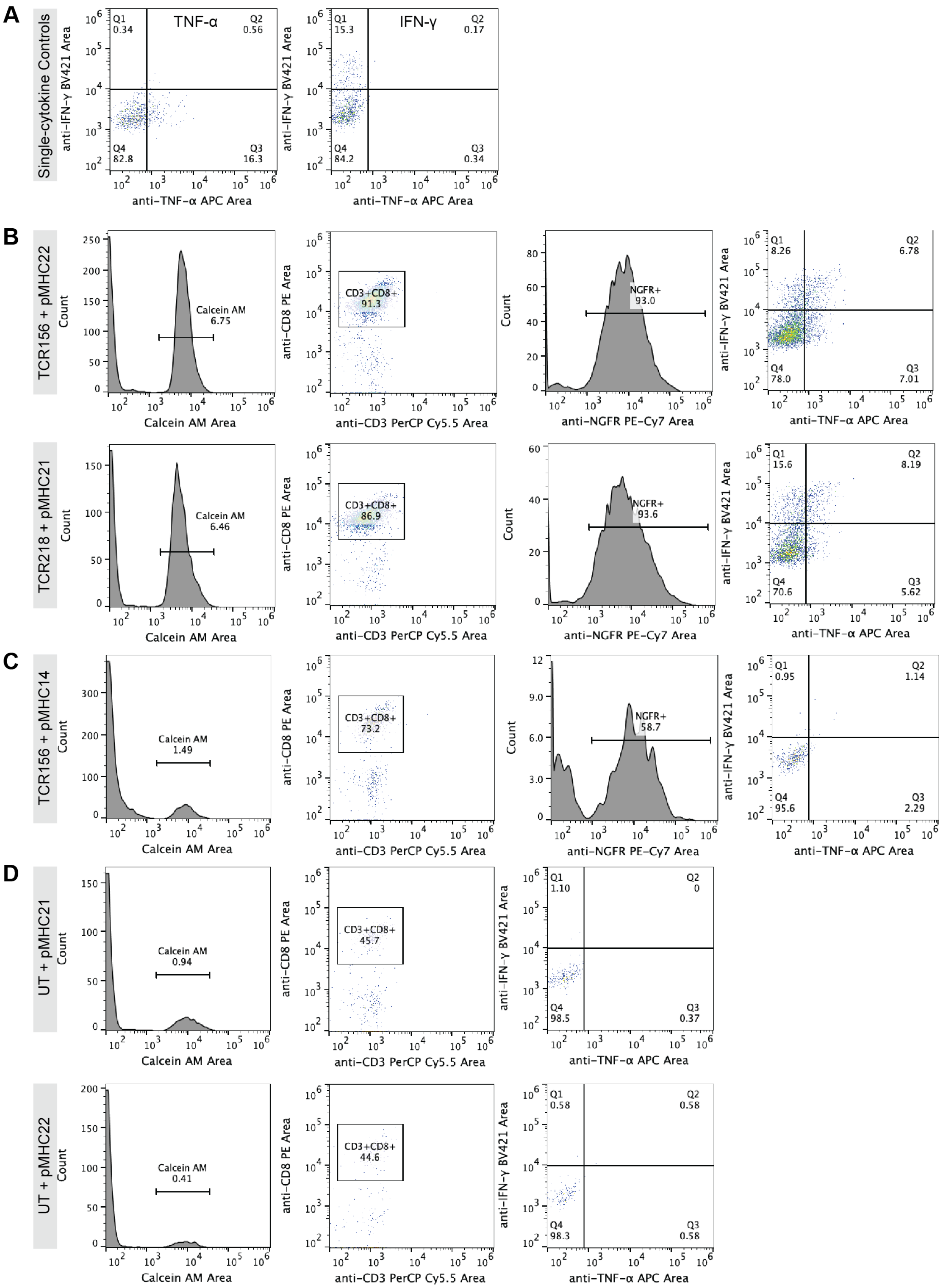


Figure S9: Detailed FACS analysis and gating strategy for multiplexed secretion-based profiling of PAP-specific T cells. (A) Flow cytometry dot plots with gating defining populations positive for IFN-γ and TNF-α secretion on nanovials with single cytokine capture antibodies. (B) Flow cytometry plots for identifying functional antigen-specific T cells transduced with TCR218 and TCR156 loaded on HLA-A*02:01 restricted PAP21 and PAP22 pMHC labeled nanovials, respectively. IFN-γ and TNF-α secretion signals were analyzed from the calcein AM stained, CD3 and CD8 double positive population with NGFR signal above background. (C) Control flow cytometry plots for analyzing secretions from TCR156 transduced PBMCs loaded on non-cognate PAP14 pMHC labeled nanovials or from (D) untransduced PBMCs loaded on PAP21 and PAP22 pMHC labeled nanovials.

| Clonotype ID | TCR_ID | TRAV | TRAJ | CDR3A | TRBV | TRBD | TRBJ | CDR3B | Frequency | Epitope |
|---|---|---|---|---|---|---|---|---|---|---|
| clonotype0_2 | TCR1 | TRAV24 | TRAJ21 | CAFISFNKFYF | TRBV6-5 | | TRBJ1-2 | CASSPQTGTGTYGYTF | 801 | CMV1 |
| clonotype0_1 | TCR4 | TRAV24 | TRAJ21 | CAFISFNKFYF | TRBV6-5 | TRBD1 | TRBJ1-2 | CASSAQTGAAYGYTF | 801 | CMV1 |
| clonotype1_1 | TCR2 | TRAV24 | TRAJ49 | CARNTGNQFYF | TRBV6-5 | | TRBJ1-2 | CASSAQTGAAYGYTF | 308 | CMV1 |
| clonotype1_3 | TCR3 | TRAV24 | TRAJ49 | CILRDIPFDRGSTLGRLYF | TRBV6-5 | | TRBJ1-2 | CASSAQTGAAYGYTF | 308 | CMV1 |
| clonotype1_2 | TCR5 | TRAV26-2 | TRAJ49 | CARNTGNQFYF | TRBV6-5 | TRBD1 | TRBJ1-2 | CASSPQTGTGTYGYTF | 308 | CMV1 |
| clonotype1_4 | TCR6 | TRAV26-2 | TRAJ49 | CILRDIPFDRGSTLGRLYF | TRBV6-5 | TRBD1 | TRBJ1-2 | CASSPQTGTGTYGYTF | 308 | CMV1 |
| clonotype4 | TCR8 | TRAV3 | TRAJ31 | CAVRDLSARLMF | TRBV12-4 | | TRBJ2-7 | CASSSVNEQYF | 41 | CMV1 |
| clonotype5 | TCR9 | TRAV17 | TRAJ52 | CAMWTSYGKLTF | TRBV7-3 | | TRBJ2-2 | CASSLEVTGELFF | 35 | CMV1 |
| clonotype11_1 | TCR12 | TRAV5 | TRAJ36 | CAERIQTGANNLFF | TRBV13 | | TRBJ1-1 | CASSLGGGGVTEAFF | 14 | CMV1 |
| clonotype11_3 | TCR13 | TRAV13-2 | TRAJ8 | CAERIQTGANNLFF | TRBV24-1 | | TRBJ1-1 | CATTYGRMNTEAFF | 14 | CMV1 |
| clonotype11_2 | TCR14 | TRAV5 | TRAJ36 | CAVFNTGFQKLVF | TRBV13 | | TRBJ1-1 | CASSLGGGGVTEAFF | 14 | CMV1 |
| clonotype11_4 | TCR15 | TRAV13-2 | TRAJ8 | CAVFNTGFQKLVF | TRBV24-1 | | TRBJ1-1 | CATTYGRMNTEAFF | 14 | CMV1 |
| clonotype9 | TCR17 | TRAV24 | TRAJ49 | CALGYGNQFYF | TRBV27 | TRBD1 | TRBJ1-2 | CASSLLATGGNGYTF | 10 | CMV1 |
| clonotype12_1 | TCR21 | TRAV24 | TRAJ49 | CARNTGNQFYF | TRBV6-5 | TRBD1 | TRBJ1-2 | CASSPQTGTGTYGYTF | 7 | CMV1 |
| clonotype12_2 | TCR22 | TRAV24 | TRAJ49 | CARNTGNQFYF | TRBV6-5 | | TRBJ1-2 | CASSRQTGSIYGYTF | 7 | CMV1 |
| clonotype14_1 | TCR23 | TRAV3 | TRAJ31 | CAVRDISARLMF | TRBV12-4 | | TRBJ1-1 | CASSSVTEAFF | 6 | CMV1 |
| clonotype14_2 | TCR24 | TRAV10 | TRAJ45 | CVVIVMYSGGGADGLTF | TRBV12-4 | | TRBJ1-1 | CASSSVTEAFF | 6 | CMV1 |
| clonotype20 | TCR25 | TRAV8-3 | TRAJ43 | CAVAPSNDMRF | TRBV27 | | TRBJ2-1 | CASSLVSGATYNEQFF | 5 | CMV1 |
| clonotype26 | TCR34 | TRAV5 | TRAJ42 | CAEIPNYGGSQGNLIF | TRBV12-4 | | TRBJ1-2 | CASSLVGGRYGYTF | 2 | CMV1 |
| clonotype51 | TCR35 | TRAV21 | TRAJ49 | CAANTGNQFYF | TRBV7-8 | | TRBJ2-3 | CASSLMFTGVPQDTQYF | 2 | CMV1 |
| clonotype36 | TCR38 | TRAV24 | TRAJ49 | CARNTGNQFYF | TRBV6-5 | | TRBJ1-2 | CASSPTTGTAYGYTF | 2 | CMV1 |
| clonotype3 | TCR7 | TRAV5 | TRAJ37 | CAESIGKLIF | TRBV29-1 | TRBD1 | TRBJ1-4 | CSVGHGGTNEKLFF | 49 | EBV |
| clonotype6_1 | TCR10 | TRAV5 | TRAJ3 | CAEYSSASKIIF | TRBV14 | | TRBJ2-1 | CASSQSPGGTQFF | 26 | EBV |
| clonotype6_2 | TCR11 | TRAV4 | TRAJ16 | CLVGDGKSDGQKLLF | TRBV14 | | TRBJ2-1 | CASSQSPGGTQFF | 26 | EBV |
| clonotype7 | TCR16 | TRAV5 | TRAJ23 | CAESIGKLIF | TRBV29-1 | TRBD1 | TRBJ1-4 | CSVGQGGTNEKLFF | 10 | EBV |
| clonotype13_1 | TCR18 | TRAV5 | TRAJ31 | CAEDNNARLMF | TRBV20-1 | TRBD1 | TRBJ1-2 | CSARDGTGNGYTF | 8 | EBV |
| clonotype13_2 | TCR19 | TRAV13-1 | TRAJ26 | CAVYGGQNFVF | TRBV20-1 | TRBD1 | TRBJ1-2 | CSARDGTGNGYTF | 8 | EBV |
| clonotype10 | TCR20 | TRAV5 | TRAJ3 | CAEYSSASKIIF | TRBV14 | | TRBJ2-3 | CASSQSPGGTQYF | 7 | EBV |
| clonotype22 | TCR33 | TRAV5 | TRAJ31 | CAEDSNARLMF | TRBV20-1 | TRBD1 | TRBJ1-2 | CSARDGTGNGYTF | 4 | EBV |
| clonotype18 | TCR32 | TRAV3 | TRAJ12 | CAVRDTRDSSYKLIF | TRBV10-3 | | TRBJ2-7 | CAISEDITVAPEQYF | 3 | EBV |
| clonotype45 | TCR36 | TRAV1-2 | TRAJ10 | CAVDILTGGGNKLTF | TRBV27 | | TRBJ1-5 | CASGPYEGNQPQHF | 2 | EBV |
| clonotype35 | TCR37 | TRAV29 | TRAJ42 | CAAGGSQGNLIF | TRBV27 | | TRBJ1-2 | CASSLTGTFRGYTF | 2 | EBV |

Figure S10: Detailed list of clonotypes recovered from nanovials (frequency >=2) with corresponding V(D)J αβ genes, CDR3 αβ sequences, frequency of clonotype and epitope information. Sequences colored in pink represent those found in previously reported studies.

**Supplementary Note**

From fluorescence microscopy images, we observed two distinct fluorescence patterns, fluorescence spread across the nanovial cavity, presumably from secreted cytokines and fluorescence associated with cells on nanovials (without signal on the nanovial). We developed an approach to use the fluorescence peak shape to distinguish between nanovial and non-specific cell staining. From fluorescence images of T cells secreting on nanovials, we plotted the fluorescence intensity profile across the cavity diameter using MATLAB and calculated the maximum intensity (height), the area under the intensity curve (area), and the ratio between the area and height (fig. SN.B). Nanovials with both spatially spread secretion signal on their cavities and localized labels bound to the surfaces of adhered cells were found to have a similar range of fluorescence peak height values. However, the area over height measurement was distinctly higher for the nanovials with secretion signal. This information may be used as a distinguishing feature in flow cytometry, as analogous fluorescent pulses are generated when nanovials pass through the excitation laser beam spot (fig. SN.A). The height of the flow cytometry pulse is determined by the maximum fluorescence intensity of the nanovial and the area integrates the intensity emitted over the entire transit event through the laser spot. Accordingly, our nanovials with spatially-distributed secretion signals are expected to produce higher fluorescence area signals for a given fluorescence intensity (height) compared to nanovials with cells bound to labels. We analyzed our samples based on a combination of fluorescence peak area and peak height signals (area vs. height plot) and observed two populations, where one population had higher area signal as compared to the other population with similar height values. When we sorted nanovials with larger ratios of area/height (2.06% high secretion and 2.77% low secretion gates), the recovered nanovials had higher secretion signals, while sorted events in the lower area/height (A/H) region (2.61% label binding to cell gate) corresponded to nanovials with label bound to cells (fig. SN.C). Using this area vs. height metric, we were able to sort populations of cells with secreted cytokine signal only (A/H > 3), completely differentiating secretion signal on nanovials from signal solely from cell surface binding or intracellular staining of permeabilized or dead cells (fig. SN.D). The percent of the cell population with label bound was consistent across all three cytokines (~2.6% of the total analyzed events).

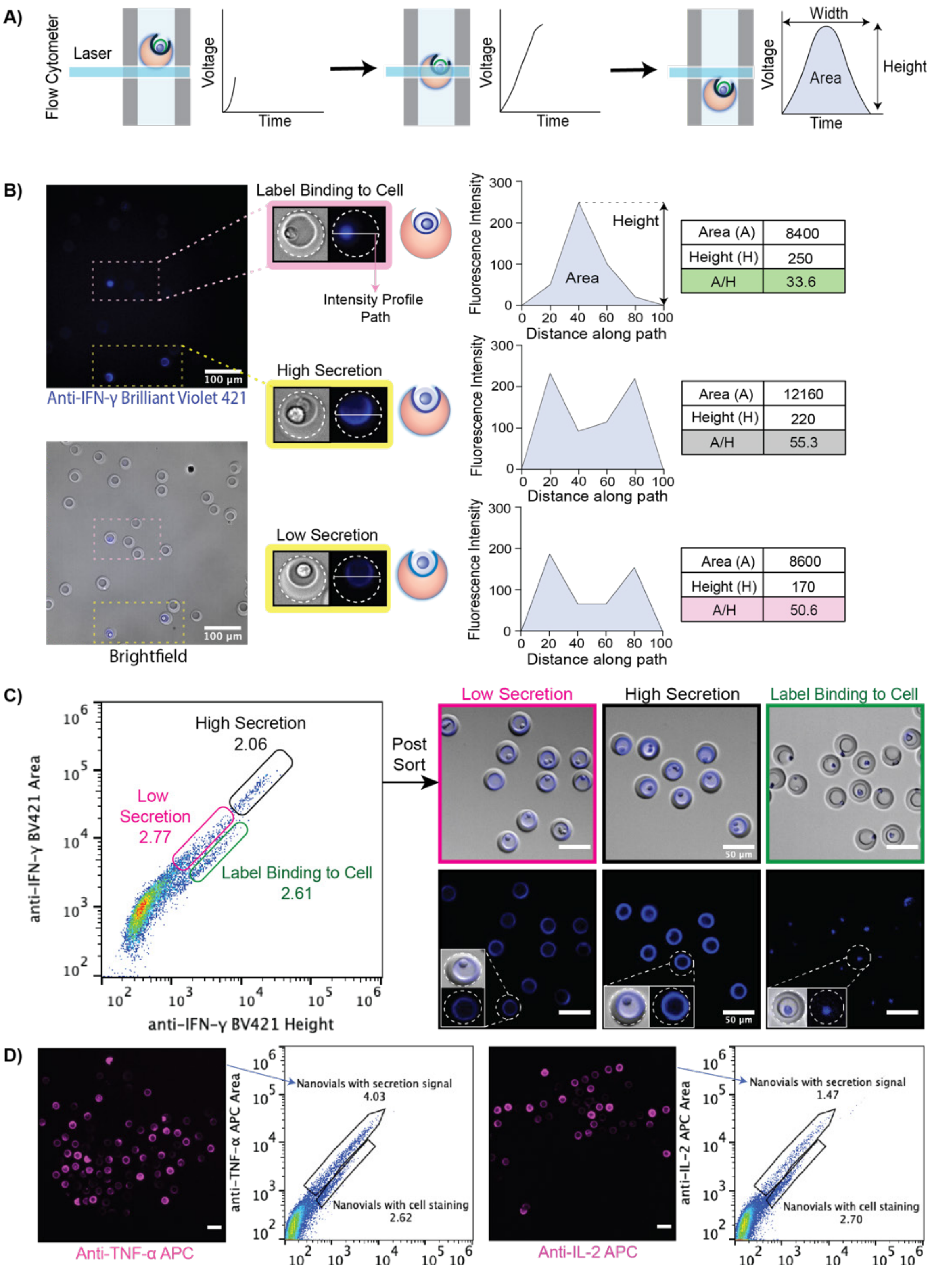


Figure SN: Using the fluorescence peak area and height to measure single-cell secretions on nanovials. A) Overview schematic of the nanovial fluorescence peak shape obtained when a nanovial transits a laser spot in a flow cytometer. B) Fluorescence microscopy images of pre-sort nanovials with cells showing two distinct fluorescence patterns, with dotted lines in insets outlining the nanovial boundaries: fluorescence spread across the nanovial cavity from secreted cytokines or fluorescence associated with cells on nanovials without signal across the cavity area. The fluorescence intensity profile was computed across the cavity of each image and the maximum intensity (height, H), area under the intensity curve (area, A), and the ratio between the area and height were calculated. Scale bars represent 100 μm. C) Isolation of T cells on nanovials with secretion signal. Three different gates were used to differentiate spatially-extended IFN-γ secretion signals on nanovials (pink and black secretion gates) from signal solely from non-specific cell surface binding (green label binding to cell gate). Sorted cells on nanovials have different distribution of fluorescence signal, as shown in the images. Scale bars represent 50 μm. D) Nanovials with each cytokine (TNF-α or IL-2) secretion signal were sorted using area vs. height metrics. The ability to isolate on-nanovial cytokine staining was consistent across different cytokines as shown in fluorescence microscopy images. Scale bars represent 50 μm.
